# A framework for the development of a global standardised marine taxon reference image database (SMarTaR-ID) to support image-based analyses

**DOI:** 10.1101/670786

**Authors:** Kerry L. Howell, Jaime S. Davies, A. Louise Allcock, Andreia Braga-Henriques, Pål Buhl-Mortensen, Marina Carreiro-Silva, Carlos Dominguez-Carrió, Jennifer M. Durden, Nicola L. Foster, Chloe A. Game, Becky Hitchin, Tammy Horton, Brett Hosking, Daniel O. B. Jones, Christopher Mah, Claire Laguionie Marchais, Lenaick Menot, Telmo Morato, Tabitha R. R. Pearman, Nils Piechaud, Rebecca E. Ross, Henry A. Ruhl, Hanieh Saeedi, Paris V. Stefanoudis, Gerald H. Taranto, Michael B Thompson, James R. Taylor, Paul Tyler, Johanne Vad, Lissette Victorero, Rui P. Vieira, Lucy C. Woodall, Joana R. Xavier, Daniel Wagner

## Abstract

Video and image data are regularly used in the field of benthic ecology to document biodiversity. However, their use is subject to a number of challenges, principally the identification of taxa within the images without associated physical specimens. The challenge of applying traditional taxonomic keys to the identification of fauna from images has led to the development of personal, group, or institution level reference image catalogues of operational taxonomic units (OTUs) or morphospecies. Lack of standardisation among these reference catalogues has led to problems with observer bias and the inability to combine datasets across studies. In addition, lack of a common reference standard is stifling efforts in the application of artificial intelligence to taxon identification. Using the North Atlantic deep sea as a case study, we propose a database structure to facilitate standardisation of morphospecies image catalogues between research groups and support future use in multiple front-end applications. We also propose a framework for coordination of international efforts to develop reference guides for the identification of marine species from images. The proposed structure follows the Darwin Core standard to allow integration with existing databases. We suggest a management framework where high-level taxonomic groups are curated by a regional team, consisting of both end users and taxonomic experts. We identify a mechanism by which overall quality of data within a common reference guide could be raised over the next decade. Finally, we discuss the role of a common reference standard in advancing marine ecology and supporting sustainable use of this ecosystem.

## 1. Introduction

There is a long history of using images in marine ecological studies. The first underwater photograph was taken in 1856 in UK seas [1] but it took until 1893, on the sunlit Mediterranean seabed, for the first clear images to be produced [2]. Following this, the use of underwater photography became widespread in shallow seas, opening up this environment to a wider public (e.g. [3]). The first deep-sea photograph was taken from the porthole of a bathysphere in the early 1930s [4] and shortly after, the first self-contained deep-sea photographic systems were developed in the 1940s at the Woods Hole Oceanographic Institution [5, 6]. Whilst there were many good deep-sea photographs available between this time and the early 1970s [7, 8], few biologists studied them, as often no corresponding samples of animals were taken, making identification difficult [9]. The notable exceptions to this [9,10, 11, 12, 13, 14] paved the way for photography to become established as an important tool for the study of deep-water environments [15, 16, 17, 18, 19]. Today, with the routine use of seafloor cameras, towed camera platforms, remotely operated and autonomous underwater vehicles (ROVs and AUVs), photographic assessment of marine fauna and faunal assemblages is a vital tool for research used by both scientists and industry [20, 21, 22].

Imaging is an important non-destructive tool for studying marine geology and biodiversity at a wide range of spatial scales (from millimetres to tens of km) [21, 23]. It enables a rapid assessment of wide areas while retaining valuable ecological information, such as spatial distribution and associations between organisms and with the landscape. Photographic and video assessment is particularly useful in complex terrain or sensitive areas [24, 25], where direct sampling is challenging or undesirable. Imaging is generally used to provide both qualitative and quantitative information on the marine environment (e.g. sediment type [26]; hyperbenthic (living immediately above the seafloor) and midwater organisms [27]; benthic epifauna (the organisms living on the sediment surface [24, 28, 29]); and faunal activity or behaviour (through visible life traces or video/time-lapse images [30, 31, 32]). As a non-destructive tool, imaging is also paramount in the identification of Vulnerable Marine Ecosystems (VMEs) [33, 34]. It has also been widely used to access the impact of human activities on benthic communities e.g. [35, 36] and to evaluate the distribution of marine litter in the seafloor e.g. [37, 38]. Imaging has also been applied to detecting and assessing temporal variation [22, 39]. Estimates of organism densities from seafloor imagery have proven more accurate than those obtained by physical sampling methods, such as trawling. For instance, densities derived from seafloor imagery provided a 10-50 fold increase in accuracy in comparison to trawling in the Porcupine Abyssal Plain in the North East Atlantic [40]. However, it is likely that diversity is underestimated as a result of difficulties of identification of the taxa to lower taxonomic levels from imagery [21].

The use of images to collect faunal data brings with it the challenge of identifying taxa from image data. Identification of physical specimens is usually achieved using taxonomic keys that have been developed by experts working on specific taxonomic groups. These keys are developed based on thorough study of preserved specimens, incorporating a systematic analysis of characteristic morphological features, followed by the development of a dichotomous key. While traditional taxonomic keys may be useful in the identification of some taxonomic groups from imagery (e.g. fish), many such keys rely on characteristics that are not visible in imagery (e.g. the arrangement of mesenteries in anemones, spicule shape in sponges, sclerite morphology in gorgonians, and the ossicles of holothurians).

Therefore, for many taxonomic groups the development of field guides are essential to support taxon identification from image data. Many field guides have been developed for shallow-water marine species for use by SCUBA divers. These rely heavily on image data to show form, function and details of anatomy that can be used for accurate identification e.g. [41, 42], but they are rare for depths beyond recreational SCUBA diving capability (>30 m) (hereinafter referred to as deep-water species). Good field guides are usually underpinned by a comprehensive understanding of the species pool for the region of study. For most deep-water regions, this understanding is lacking. Notable exceptions include the Monterey Canyon [43] and the soft sediment (trawlable) habitats of the North Atlantic. The lack of comprehensive field guides for deep-water marine organisms presents a significant challenge to those faced with the interpretation of image data from poorly known regions or habitats, such as seamounts, ridges, or other areas of hard and high-relief substrates that are not conducive to trawling surveys.

In the absence of a good knowledge of the taxonomy of many groups and regional field guides, a common practice in the interpretation of image data is the development of a morphospecies reference image dataset (Fig. 1) and the use of operational taxonomic unit (OTU) numbers. The OTU numbers are used in place of taxon names for organisms for which a species name has not yet been assigned owing to the lack of physical specimens to corroborate the observation [24, 43, 44, 45, 46, 47, 48, 49, 50]. These morphospecies reference image catalogues provide a permanent reference of what has been observed in the study. But perhaps more importantly, allow the user to differentiate between taxa below the lowest level of the taxonomic hierarchy to which the observed organism can be identified, using traditional taxonomic features, and thus preserve important information on biodiversity. For example, taxonomic identification of many sponge and soft coral species is impossible from image data alone, since their taxonomy is based on the arrangement, size and shape of microscopic structures in their skeletons. Thus, following traditional methods of sample analysis, all observed species would be assigned the level Porifera or Alcyonacea, resulting in a significant loss of resolution in the data. However, use of a morphospecies reference image catalogue allows the observer to assign morphologically different (and in most cases, likely taxonomically distinct) forms to a unique OTU number, which can then be assigned to the taxon (e.g. Porifera msp. 1, Porifera msp 2 etc.) if needed, thereby retaining taxonomic resolution in the data.

**Figure 1:**
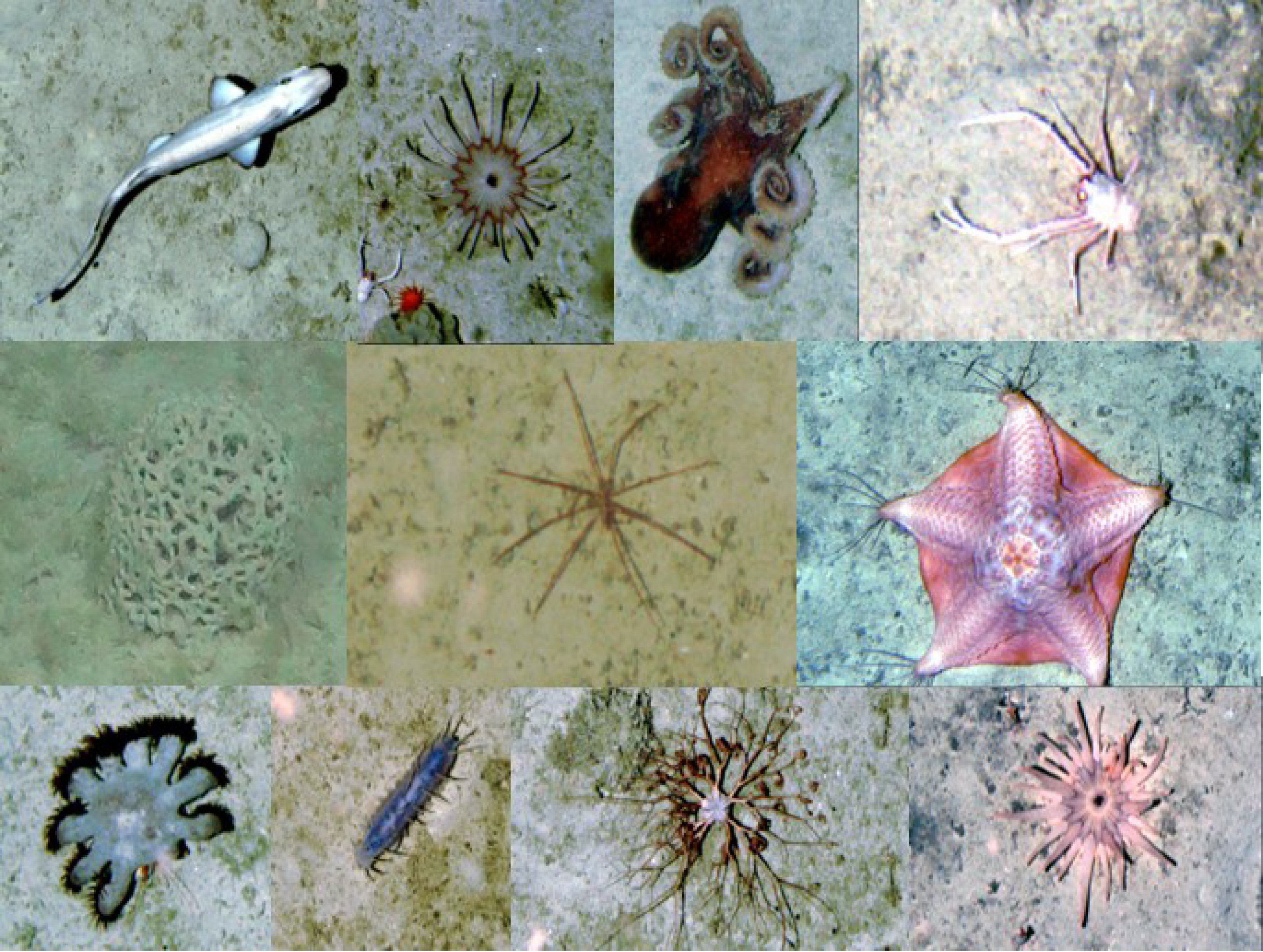
Example of a reference image catalogue where representatives of each taxa observed are cropped from an image, and assigned an OTU number that is subsequently used in image analysis in place of a standard latin name.

The problem with this approach is that each study or group uses a different naming convention for morphospecies. It then becomes impossible to compare or combine datasets between studies. Morphospecies catalogues are not usually published, making it difficult for researchers to compare data or check identifications. Comparison between research studies or industry-gathered data (for example from environmental impact assessments or site monitoring) are also impaired by this issue. In addition, both field guides and morphospecies reference image catalogues fail to document explicitly the visual characteristics used to differentiate taxa. They generally provide little more than a visual idea of what a taxon looks like. This compounds problems of observer biases that are well documented in biological sample analysis [51, 52, 53]. When identifying taxa from image data, it is necessary to use a combination of traditional taxonomic features and ecological data (e.g. depth, location, habitat, knowledge of the local species pool) to arrive at an identification. This skill in ‘field identification’ is often acquired through an ‘oral tradition’ with little in the way of formalised training materials provided to new researchers entering the field or new consultants provided with image data to analyse.

Developments in autonomous and robotic technology, and the increased use of them across different fields, are increasing the amount of image-based data that can be collected [54, 55, 56]. For example, a single 22-hour AUV mission returned over 150,000 seafloor images [40, 56]. Manual image analysis is a time-consuming process, which forms the current bottleneck in image-based ecological sampling [21, 57, 58, 59]. As a result, a number of research teams are investigating the use of artificial intelligence (AI) and computer vision (CV) as potential means to accelerate and standardise the interpretation of ecological image data [51, 52, 53, 56, 60]. The most promising of these techniques is supervised machine learning to automatically detect and classify taxa [53, 58, 61]. However, consistent interpretations by humans are initially required, providing ‘gold standard’ classifications, with as much data as possible, which can be used to train these algorithms. Moving forward, developments in AI and CV approaches that combine the use of visible morphological characteristics with deep learning, would benefit significantly from the development of a standard image-reference dataset. For those taxonomic groups in which the morphological characteristics commonly used to differentiate taxa are not discernible in images (e.g. sponges, anemones, zoanthids and plexaurid gorgonians), these types of combined approaches will first require development of novel visual multi-access keys, which themselves can only be created from a high-quality reference image dataset and skilful determination of characteristics differentiating taxa.

Table 1 provides a list of field guides and morphospecies reference image catalogues for deep-water species of the Atlantic Ocean that are currently publicly available. However, many more are un-published or inaccessible to others, and are held as a mixture of printed and electronic materials. Recently there have been attempts to make morphospecies reference image catalogues associated with specific research programmes or projects available to others (for example [43, 47, 62, 63, 64, 65, 66] to mention a few). In addition classification based approaches to this issue have also been developed [67]. While useful, this ‘piece-meal’ approach will not solve the challenges outlined above.

**Table 1.**
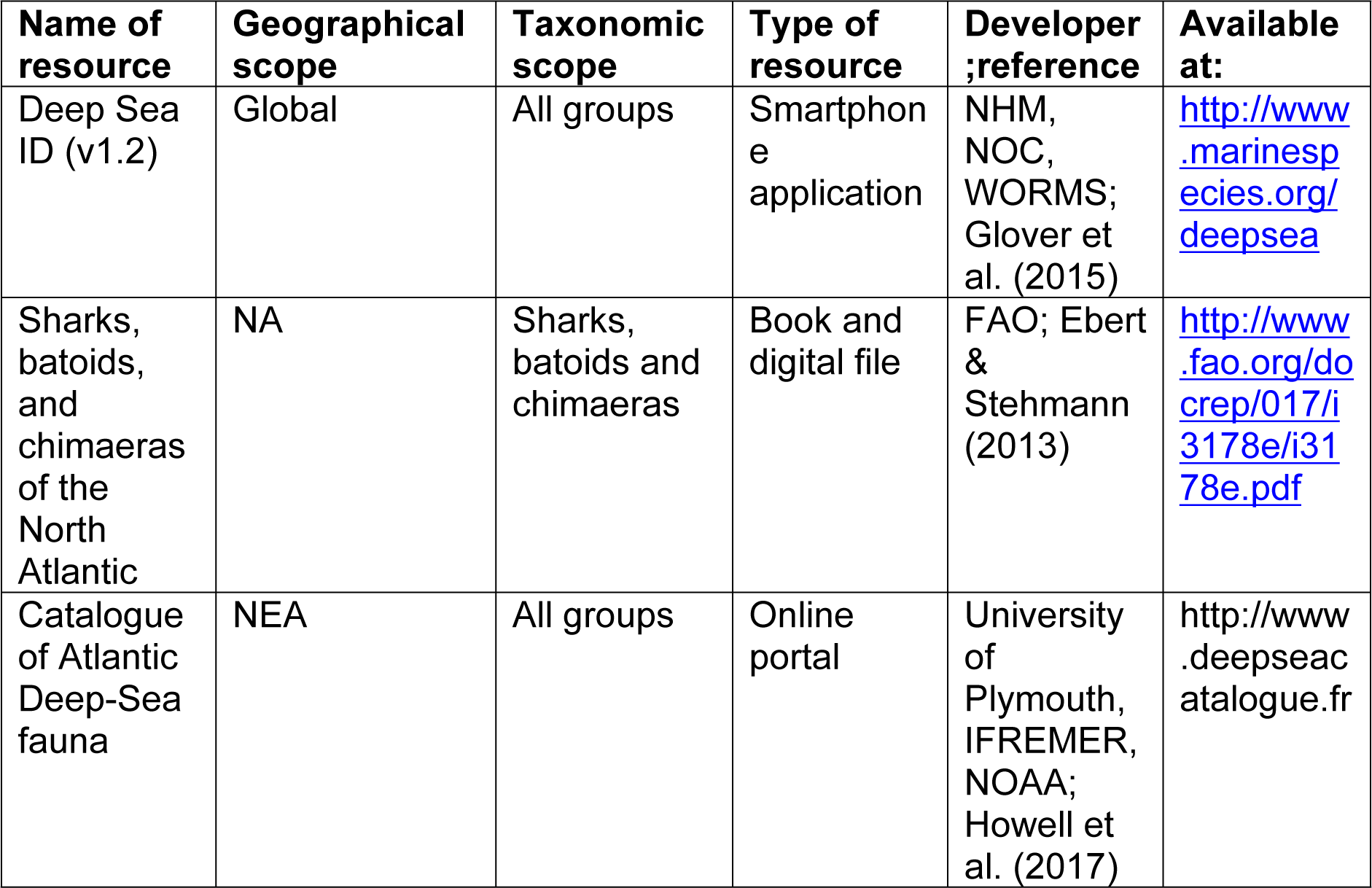
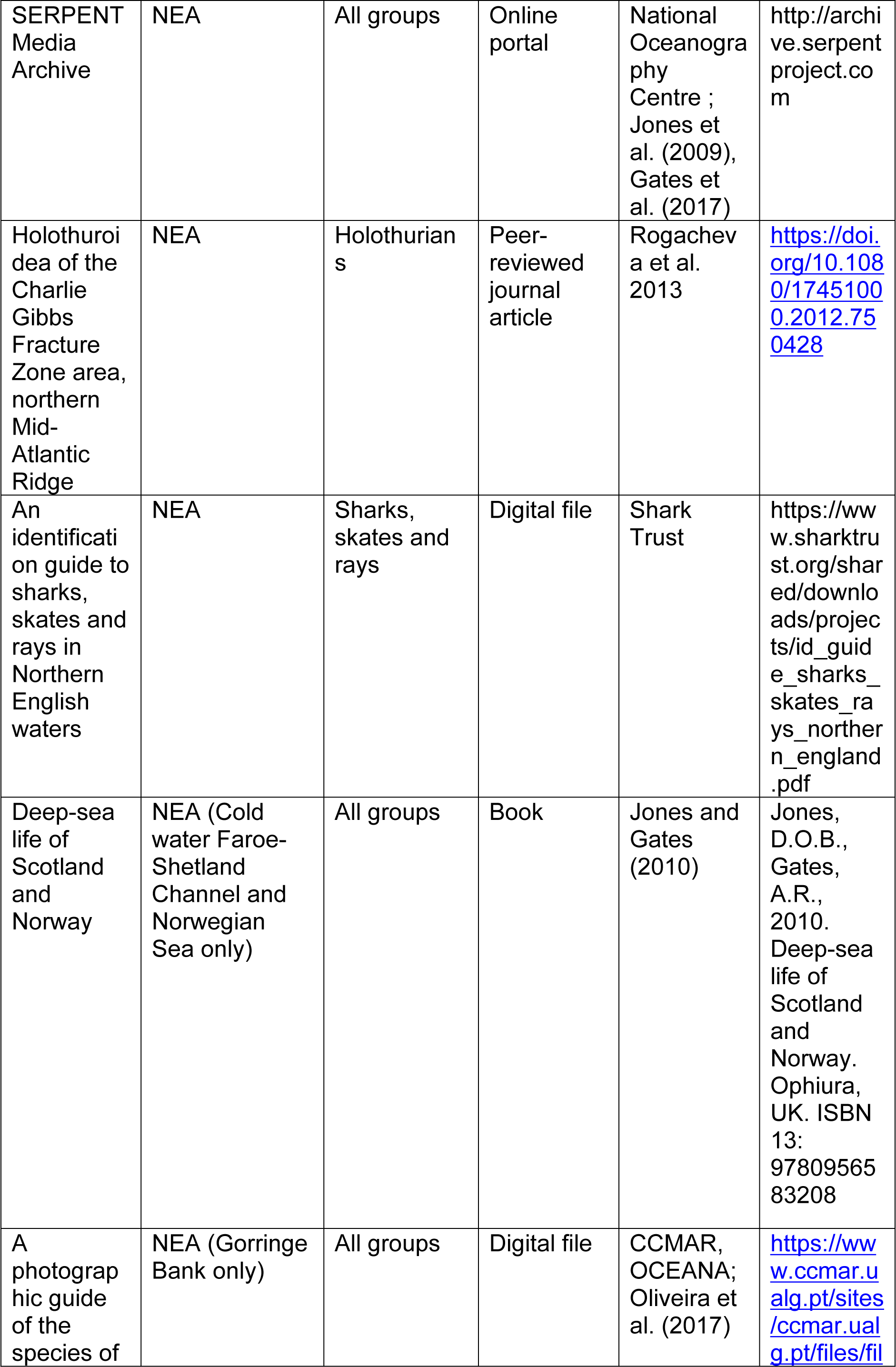
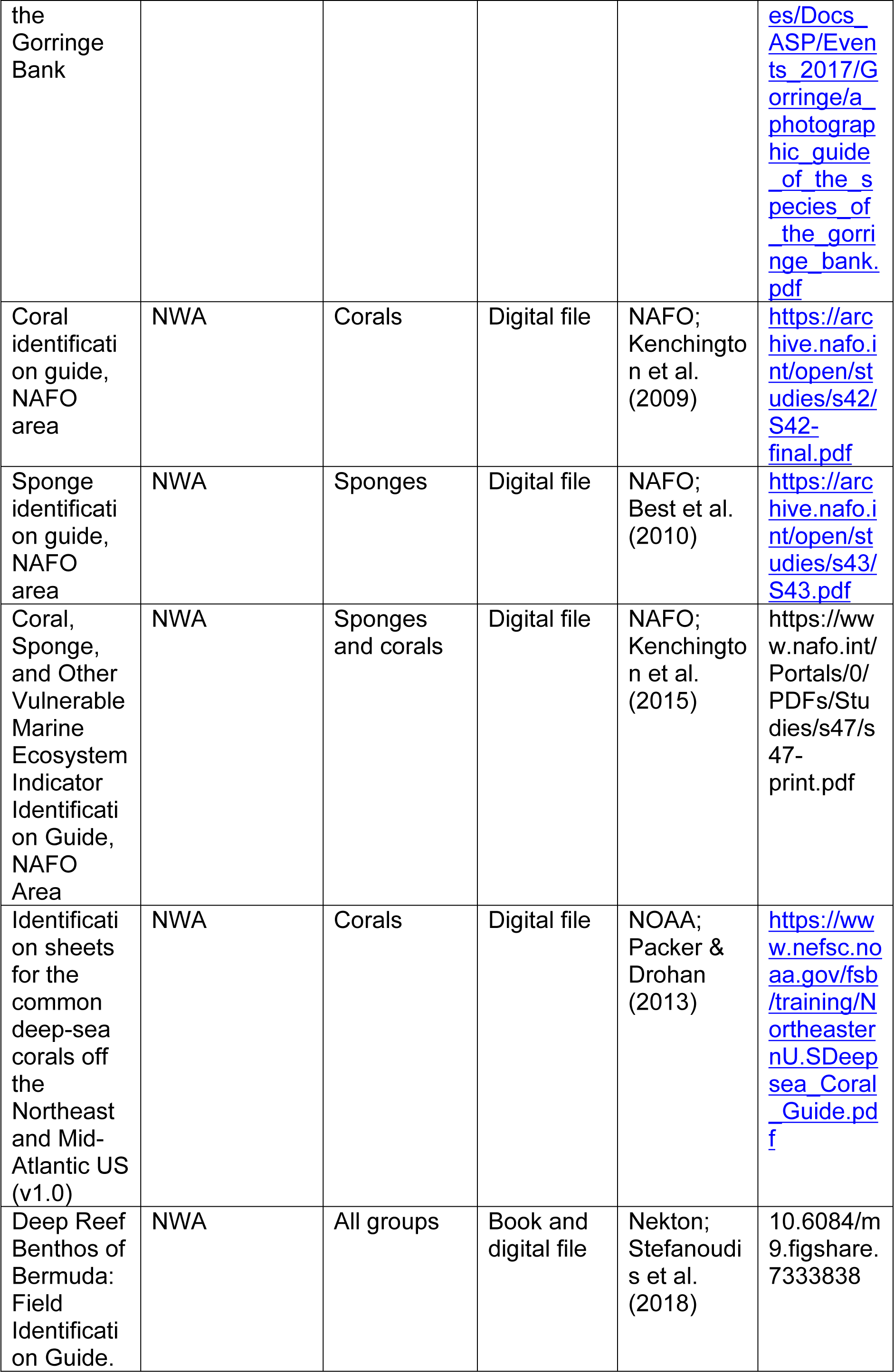
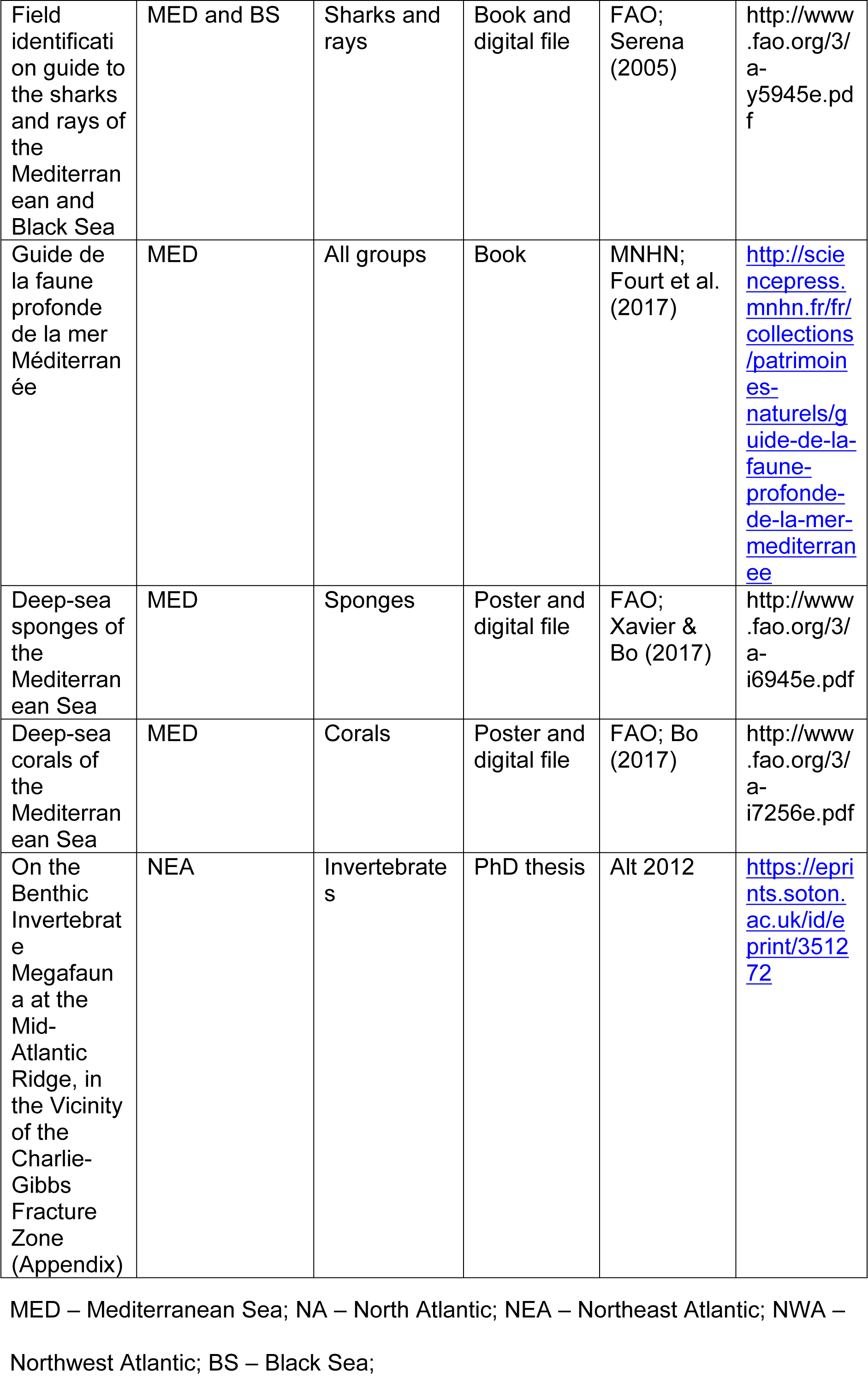
List of available image catalogues and identification guides of the deep-sea fauna off the Atlanto-Mediterranean region.

There is a clear need for the development of a standard reference guide to support the use of image-based sampling. Failure to develop appropriate tools will ultimately hinder progress in marine ecology, particularly in deep-sea marine ecology where images are frequently one of the few collected datasets. In order to improve data quality and comparability, realise the benefits of new technologies in both image data collection and interpretation, and ultimately raise standards of taxonomic identification within academia, government, and industry, we must move towards the use of standard reference guides, quality controlled and curated by experts in both taxonomy and field identification.

Our aims were to develop 1) a database structure to facilitate the standardisation (and ultimately pooling) of morphospecies reference image catalogues between individuals and groups, supporting onward use in multiple applications; and 2) a framework for coordination of international efforts to develop reference guides for the identification of deep-water species from image-based data.

## 2. Methods

The initial stages of developing the framework for the database consisted of assessing the requirements of those working with image-based data. This included the need for both online and offline databases and printable catalogues for use in making identifications at sea.

We reviewed current relevant databases and database standards. These were focused around the Darwin Core standard, the Ocean Biogeographic Information System (OBIS), and the World Register of Marine Species databases (WoRMS).

The Darwin Core is an international standard set of terms and definitions that facilitates sharing biodiversity data [68]. The Darwin Core quick reference guide (http://rs.tdwg.org/dwc/terms/), provides a comprehensive glossary of terms (standardised fields with descriptors and examples) to ensure data concerned with the occurrence of organisms, the physical existence of specimens in collections, and related environmental information can be standardised. Darwin Core forms the basis of a number of existing online open-source relevant databases (e.g. [69, 70, 71, 72]), and, thus, is the internationally agreed standard upon which further database development should be based. Darwin Core Archives (DwC-A) comprise a set of text files, including both the dataset (.csv) and a document (.xml) which describes the included files, fields, and their relationships. This offers a standard format used to describe biodiversity data and is being commonly employed to share more complex and structured datasets.

OBIS [71] was originally developed as the information management component of the Census of Marine Life (2000-2010) programme. OBIS founder, Dr. J. F. Grassle, articulated the vision of OBIS as “an online, worldwide atlas for accessing, modelling and mapping marine biological data in a multidimensional geographic context”. The OBIS database currently consists of over 55 million observations of nearly 124,000 marine species. In 2009, OBIS was adopted as a project by the International Oceanographic Data and Information Exchange (IODE) programme of the Intergovernmental Oceanographic Commission (IOC) of UNESCO. It represents an internationally important archive for species distribution data. OBIS is closely linked with WoRMS, which provides the taxonomic backbone, and geospatial data are provided by the Marine Regions database. Additional functionality includes the taxon match tool for resolving names used by other similar platforms, providing crucial quality control support for taxonomic data among the research community and biodiversity platforms [73].

WoRMS is an authoritative classification and catalogue of marine names including information on synonymy, and is curated by around 400 taxonomists globally, in accordance with best practice [72, 73, 74]. The content of WoRMS is managed by taxonomic and thematic experts, who are responsible for controlling the quality of the information contained within the database [73]. WoRMS is underpinned by the Aphia platform, which is a Microsoft Structured Query Language (MS SQL) database, containing over 400 fields spread over more than 80 related tables. This infrastructure is designed to capture taxonomic and related data and information. WoRMS is also the basis of the World Register of Deep-Sea Species (WoRDSS), which, through its app, Deep Sea ID [75], represents one of the few existing image-based deep-sea species guides (but see Table 1).

The Marine Regions database [76] provides a standard, relational list of geographic names, coupled with information and maps of the geographic location of these features. All geographic objects of the Marine Regions database have a unique ID, called the Marine Regions Geographic Identifier (MRGID). The different geographic objects are determined by a placetype and coordinates. While the coordinates are represented as different vector data types being a point, a line or a polygon, a placetype provides contextual information to the geographic objects, for example a sea, a bay, a ridge, a sandbank or an undersea trench.

Following the initial review of relevant databases and database standards, a strawman database architecture, to facilitate the standardisation of morphospecies reference image catalogues between individuals / groups, was proposed and circulated to an international team of end users, database specialists and programmers. An international workshop funded by the Deep-Sea Biology Society was held at Plymouth University, UK, on the 4^th^-5^th^ December 2017, where the draft structure was reviewed and refined. The workshop consisted of a cross section of attendees including major dataset holders, computer scientists, taxonomists, benthic ecologists, and representatives from WoRMS / WoRDSS. Following the workshop, the refined structure was tested by both workshop participants and members of the wider community, who input their existing morphospecies reference image catalogues into the new database structure. This resulted in further minor changes and the development of the final database structure.

Workshop participants also considered how to coordinate international efforts to develop reference guides to the identification of deep-water species from images. The following questions were considered by the workshop attendees, how can we: 1) merge existing published and unpublished catalogues? 2) manage new submissions to a merged catalogue? 3) improve the scope and quality of the image data within a merged catalogue? and 4) improve and classify the quality of identification from images?

## 3. Results

### 3.1. End product needs

Workshop participants, and specifically those engaged in image-based analysis, felt the most critical tools urgently required to support their work were *in-situ* photo-guides in book format (hard copy or e-book), a standard reference morphospecies taxonomic tree (or annotation scheme) that can be imported into different annotation software, and on-line user-friendly image reference catalogues that include information on characteristics used to classify animals as belonging to a particular OTU. The database structure must therefore be such that these end-use products can be easily created from the database by a query using purpose-built web-accessible software as part of future developments.

### 3.2. Database structure

The final database structure consists of two tables that contain Darwin Core fields together with additional fields for which no Darwin Core equivalent could be established. Table 2a is the OTU table. It documents the OTU, and primarily draws fields from the Darwin Core classes “Taxon” and “Identification”. Table 2b is the image table. It documents the individual image file and draws fields from multiple Darwin Core classes, including “Occurrence”, “Identification”, “Event”, “Location”, “Record-level”, and “Organism”. The two tables are related via the “OTU” field. This structure allows a single OTU (one entry into table 2a) to be related to multiple example images of the OTU (many entries in table 2b).

**Table 2.**
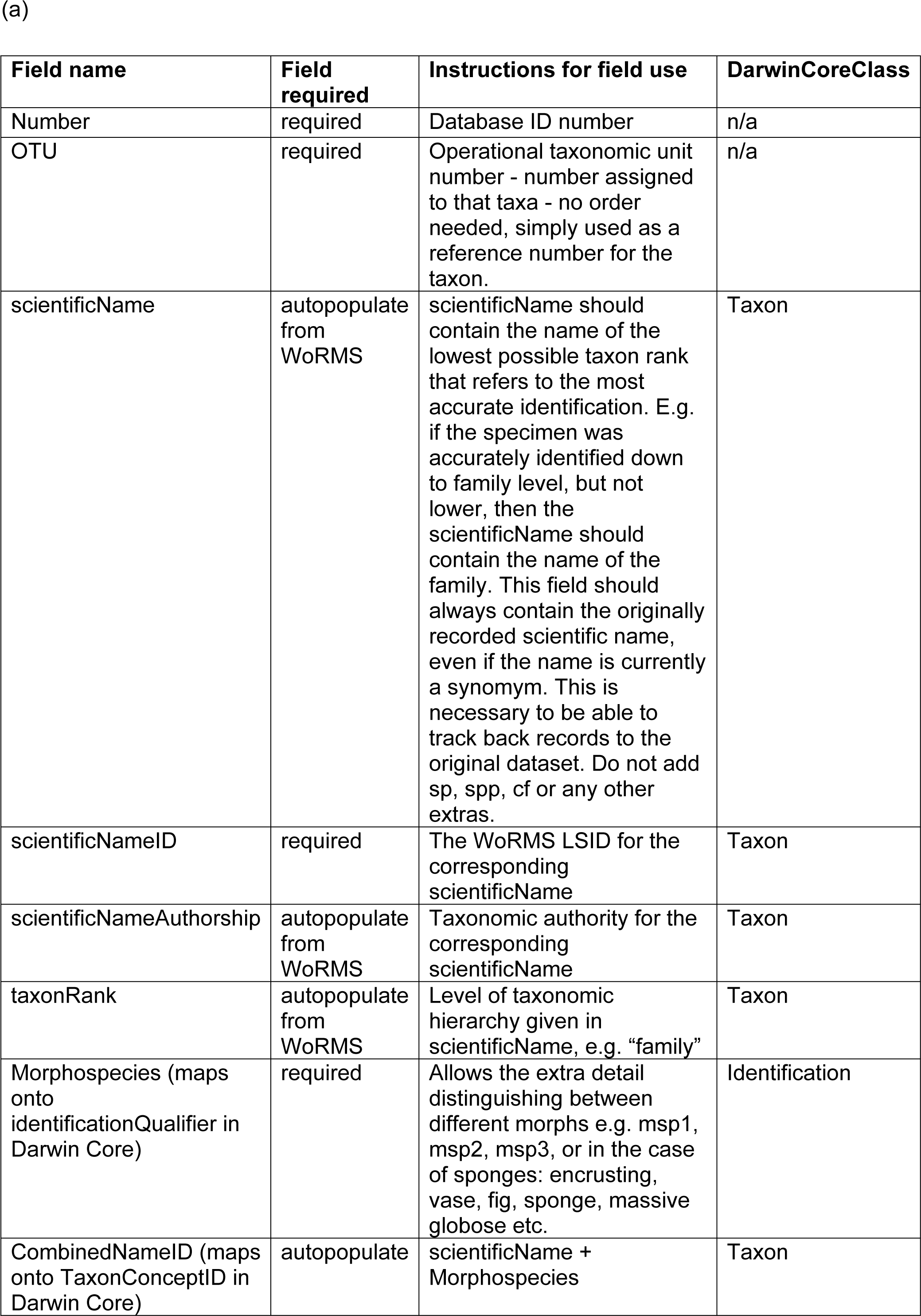
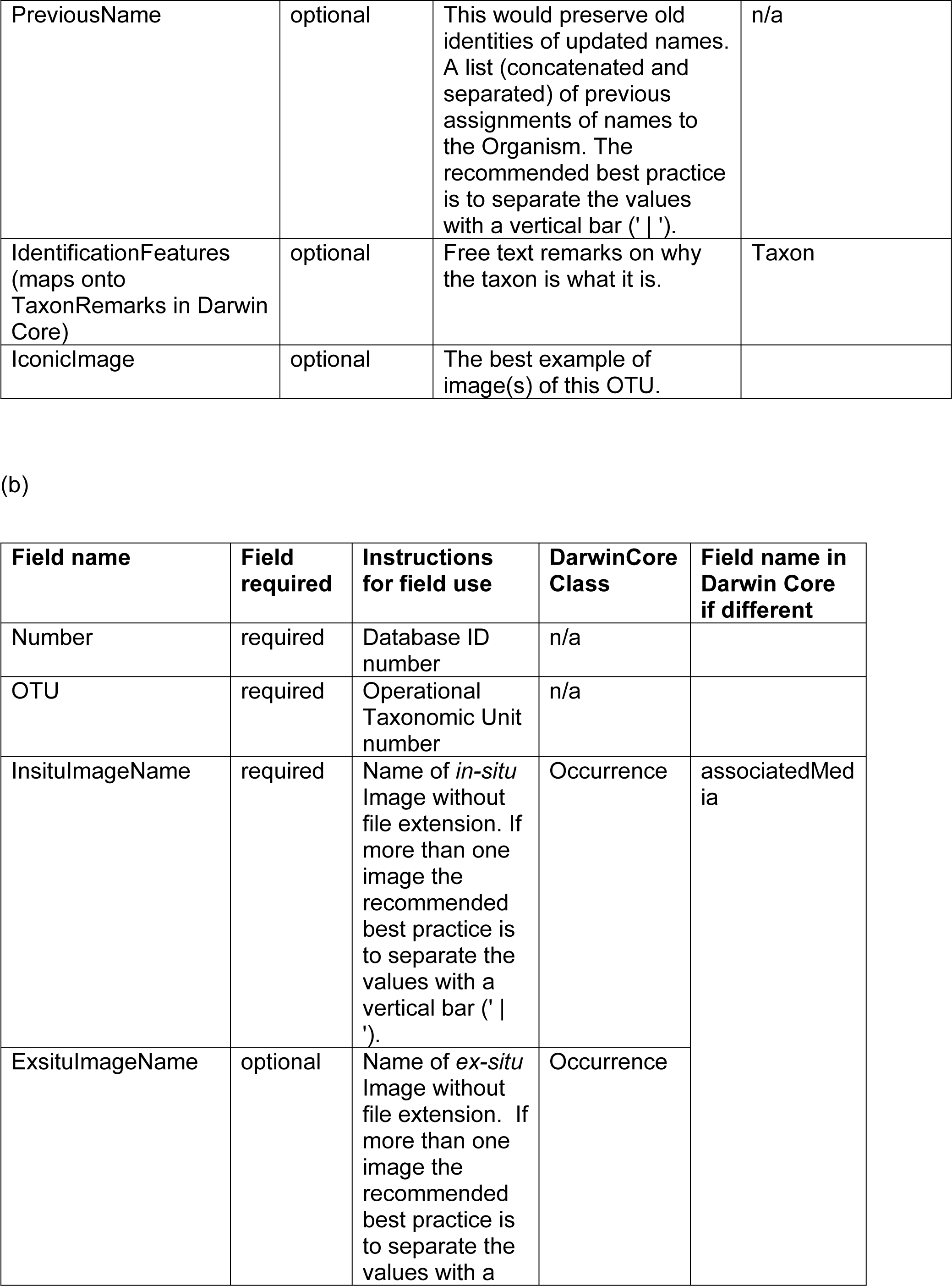
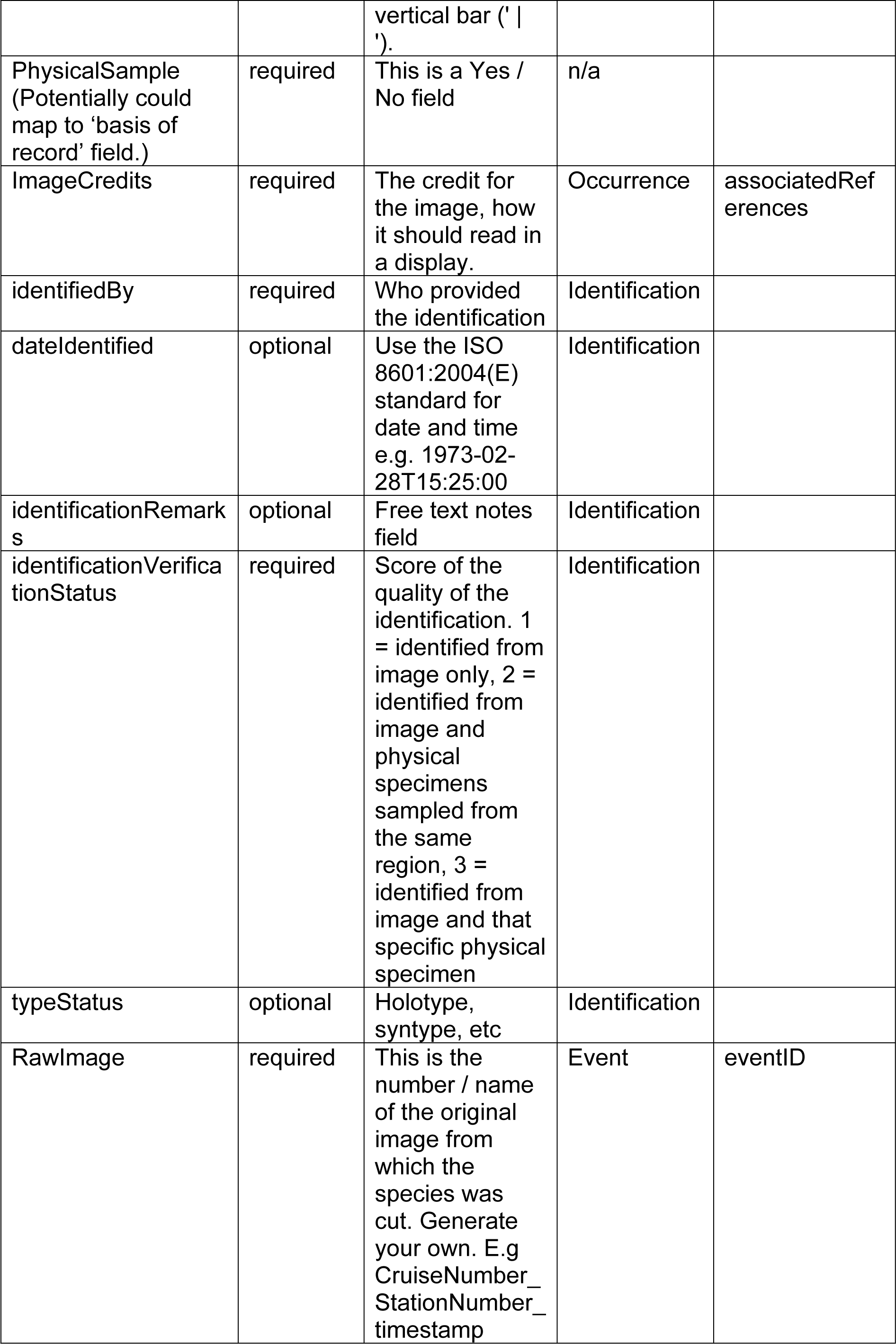
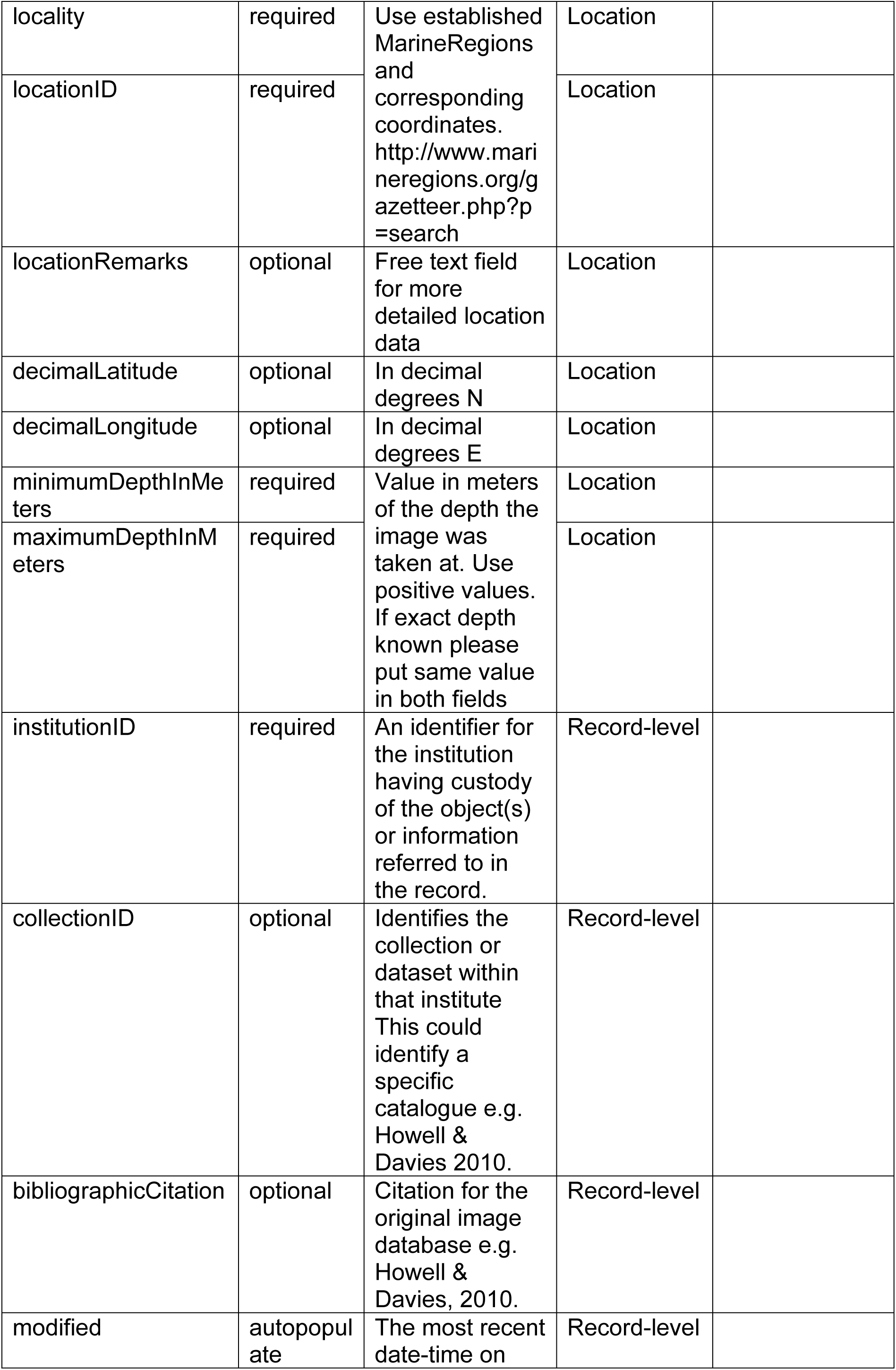
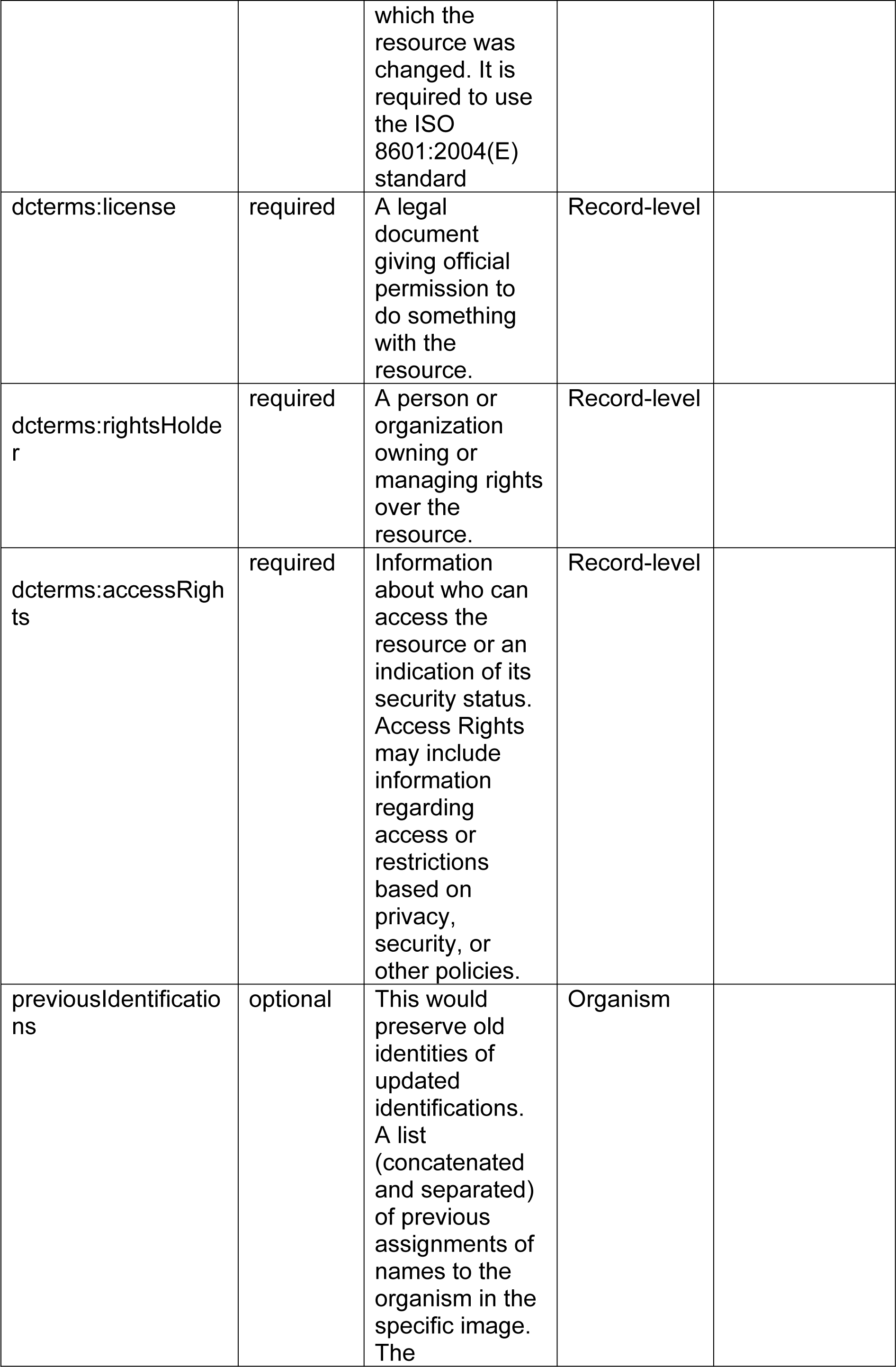
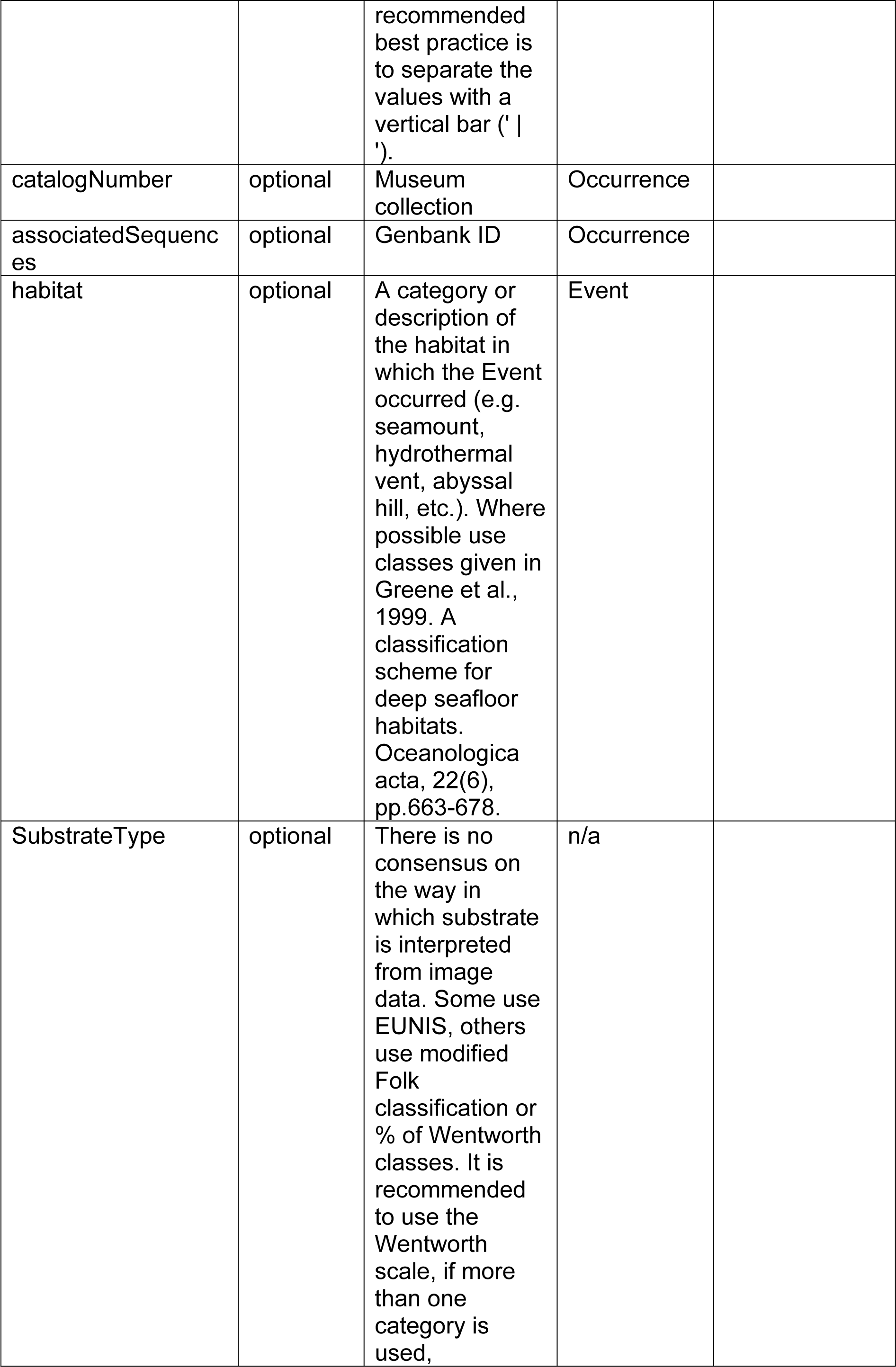
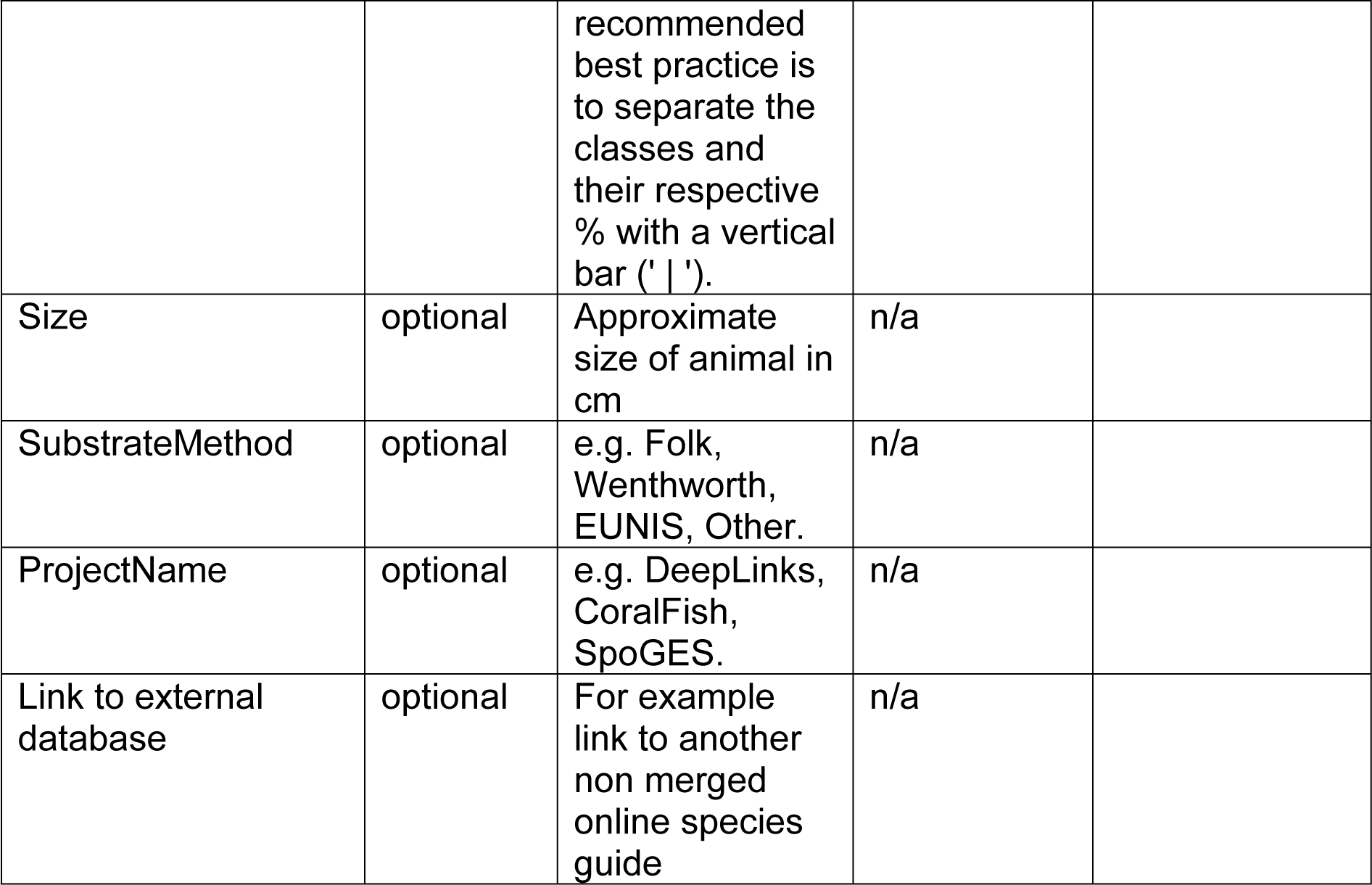
Final database structure consists of two tables related via the OTU field, the Operational Taxonomic Unit table (a), and the Image table (b)

The OTU table (Table 2a) consists of a simple index field “Number”, the inclusion of which is standard practice in database tables. The “OTU” field is a unique number given to this taxon and is initially assigned by the user. The subsequent four fields: “scientificName”, “scientificNameID”, “scientificNameAuthorship”, “taxonRank”, provide the link to the WoRMS database. The link is via the “scientificNameID” field, which requires the user to input the appropriate Life Science Identifier (LSID) for the OTU drawn from the WoRMS database. Each taxon in WoRMS receives a unique and persistent identifier, known as the AphiaID. This AphiaID can be expanded to a LSID. WoRMS has implemented LSIDs for all its taxonomic names and they are displayed on each taxon page. The LSID integrates the AphaID and so is the preferred option, of the two possible fields, to use as a link. The appropriate LSID for an OTU is the lowest formal taxonomic rank that can be assigned to an image. For some taxa, this may be at the species level; however, for many image-based identifications it will be at a higher taxonomic level, such as Family, Class or Phylum level. Use of the LSID field ensures that the OTU can be linked to standard taxonomic nomenclature and the related taxonomic hierarchy. Using this LSID, the other three fields within the database (“scientificName”, “scientificNameAuthorship”, “taxonRank”) can be auto-populated from WoRMS.

The “Morphospecies” field is equivalent to the “identificationQualifier” field in Darwin Core and allows the input of extra details distinguishing between different morphotypes; for example, Brisingidae msp1, or in the case of sponges, Porifera encrusting msp1, Porifera branching msp1. Thus, entries into this field will be of the form msp1, msp2, encrusting msp1, branching msp1, etc. The “CombinedNameID” field is then autopopulated by adding the “scientificName” and “Morphospecies” fields to give, for example, Brisingidae msp1, Porifera branching msp1. The “CombinedNameID” field can be mapped onto the “taxonconceptID” Darwin Core field. A recommended best practice for the standardisation of entries to the “identificationQualifier” field, specifically related to nomenclatural qualifiers used in image analyses is now in preparation. The “PreviousName” field is not intended to document recombinations of taxonomic nomenclature as this is captured and managed in WoRMS [74]. Rather, this field is to capture changes to the assigned identity of the OTU. For example, where Brisingidae msp1 was later confidently identified to a lower taxonomic level (e.g. *Brisinga* msp4). This field would capture its former “CombinedNameID”. The inclusion of the “IdentificationFeatures” free text field is intended to provide insight into the visual characteristics that observers are using to distinguish between morphospecies. It is hoped that over time this field will provide the material to start developing novel visual keys. The “IdentificationFeatures” free text field may map onto the Darwin Core “TaxonRemarks” field. Finally, the “IconicImage” field is used to identify the best example image of the OTU present in the database. This field determines the image that is supplied back to the WoRMS database for use on the appropriate taxon page.

The Image table (Table 2b) also has a simple index field “Number”, followed by the “OTU” field, which provides the relational link to the OTU table (Table 2a). The fields “InsituImageName” and “ExsituImageName” provide the relational link to the images that make up the morphospecies reference image catalogue, and are the name of the image file minus the file extension (e.g. IMG10542 not IMG10542.jpg). The “ImageCredits” field ensures the owners of the image are identified. We discussed at length how best to include *in-situ* and associated *ex-situ* images. While a strong argument was made around the need for good *ex-situ* images of taxa for use in developing guides for fisheries observer monitoring of bycatch, the group felt the focus of the database should be to provide a tool for the interpretation of *in-situ* image and video data. Therefore, *ex-situ* images should only be included in the database together with an accompanying *in-situ* image of the same individual. As a result, the “InsituImageName” field is required, while the “ExsituImageName” is optional. Where a physical sample has also been taken, this should be indicated in the “PhysicalSample” field as a simple yes or no. If this physical sample has been archived in a museum collection, the catalogue number should be included in the “catalogNumber” field. If it has been identified using molecular techniques, the Genbank ID should be included in the “associatedSequences” field.

The fields pertaining to the Darwin Core class “Identification” concern the identification of the individual in the image, and are self-explanatory (“identifiedBy”, “dateIdentified”, “identificationRemarks”). The “identificationVerificationStatus” field is the indicator of the quality of the identification provided. Durden et al. [21] suggest three categories of image quality: 1 = Unconfirmed: the status of the organism is uncertain, pending field collection and further taxonomic investigation, or the description and naming of a new species, 2 = Provisional: the organism is very likely this species/taxon based on investigation (literature search, consultation with outside taxonomic experts, 3 = Certain: the organism has been collected and has been definitively identified by a taxonomic expert. We have modified these categories as follows: 1 = identified from image only, 2 = identified from image and physical specimens sampled from the same region, 3 = identified from image and physical specimen of the actual individual in the image. There are often instances where an organism has been identified from an image and a specimen collected that has not yet been identified. Under these circumstances the quality score would be 1, but the existence of a specimen noted in the “PhysicalSample” field. Once a specimen is identified the quality score for the image could be changed to 2 or 3.

The fields pertaining to the Darwin Core class “Location” concern where the image was taken. We recognise that for older image data archives, exact position data may not have been recorded. However, the importance of location and depth to field identification of taxa cannot be understated. We feel it is important to ensure that the terminology used to define location is consistent with a published standard. In addition, we want to ensure that, in the future, users will be able to construct local morphospecies reference image catalogues based on selection of an area through mapping software. The Marine Regions database [76] is ideally placed to provide this geospatial standard. Its use will also ensure compatibility with OBIS such that this database can share data with OBIS and vice versa. The required fields “locality” and “locationID” provide the link to the Marine Regions database. The user must input the appropriate “locality” and “locationID” for the image drawn from the Marine Regions database. The “locationRemarks” field is an optional free text field that allows users to capture more detailed location information that is not captured by the options available in the Marine Regions database. The fields “minimumDepthInMeters”, “maximumDepthInMeters” are also required as species distributions are structured with depth [77] and this characteristic is likely to be important in the development of future field guides. The remaining fields, “decimalLatitude”, “decimalLongitude”, are optional so as to accommodate older data and / or sensitive data, for example, from industry partners.

The fields pertaining to the Darwin Core class “Record-level” focus on ownership and origin of the image. Required information includes the name of the institution that owns the image (“institutionID”), a licence document (“dcterms:license”), the name of the person / institution managing right over the image (“dcterms:rightsHolder”), and the terms of access to the image (“dcterms:accessRights”). It is anticipated that a standard licencing arrangement can be agreed to upon submission of material to the database, whereby image ownership is retained by the organisation / individual submitting but use for scientific purposes is freely granted. Use of images for commercial gain would be prohibited. There are existing licencing models for WoRDSS and these can be replicated here. Optional fields allow the identification (“collectionID”) and citation (“bibliographicCitation”) of any previously published or in-house morphospecies reference image catalogues from which the image data have been drawn. The modified field is autopopulated and is the most recent date-time on which the resource was changed.

There are just two fields that relate to the image collection event via the Darwin Core class “Event”. These are the fields “RawImage”, which is equivalent to the Darwin Core “eventID” field, and “habitat”. It is not the intention of this database to capture details of the research cruises, ROV dives, etc., on which the organism images were taken. These details are not overly important to the creation of a field guide. However, should this information be viewed as important in the future, we suggest that images are given the name of the original image from which the organism was cropped, and that this name be extended to consist of the following elements: CruiseNumber_StationNumber_timestamp_imagename. The “habitat” field is able to capture the geomorphological setting in which the organism was observed, e.g. seamount, canyon, mid-ocean ridge. We felt this information might be useful in the development of a field guide. The ideal situation would be to use standardised terms to describe these settings. We suggest the use of Greene et al. [78] as a standard reference; however, the European Nature Information System (EUNIS) [79, 80] or other classification systems may also provide a reasonable standard and the standard used could be indicated when data are submitted. One final field is drawn from the Darwin Core class “Organism” and is used to capture previous names that have been assigned to the organism in the image (“previousIdentifications”). As with the “PreviousName” field in the OTU table, this field is not used to capture taxonomic name changes, which are well recorded by WoRMS. It is used to capture changes in opinion on the identity of the organism in the image.

The remaining fields in the Image table are not Darwin Core fields but do provide additional information that is important to record. The “SubstrateType” field allows details of the substrate on which the organism was observed to be logged. Substrate is an important environmental factor that determines the distribution of species and can play a role in the field identification of taxa. As always though, it is preferable to use standard terminology to record substrate and there are many standards available. Among workshop participants, there was no consensus on methods of substrate interpretation from image data, and the terminology standards used. Some use EUNIS [79, 80], some a modified Folk [81] classification and others percentage of Wentworth [82] sediment size classes. The “SubstrateMethod” field allows the user to indicate the standard they have followed. The “Size” field, standardised to centimetres, is self-explanatory and may be useful in the future development of a field guide. The “ProjectName” field offers the opportunity to credit specific projects with provision of imagery, while the “Link to external database” field enables links to be made to source on-line morphospecies reference image catalogues.

The images are not stored within the table itself but should be provided as separate image files. Those with existing morphospecies reference image catalogues have tended to either paste images into Word or Power Point files, organise their data as Apple ibooks, or organise their images into Phylum or Class level folders. While this is useful at an individual level, and provides the end product required, it limits onward use and is not the appropriate format for a database.

### 3.3. A framework for coordination

While the database structure outlined above provides the means to archive and exchange data, the development of a unified morphospecies reference image catalogue requires a management structure to curate the database and manage new data submissions. The WoRMS database provides a model that can be adapted for use with this database. WoRMS is curated by teams who are responsible for different taxonomic groups. Each team is led by an editor who takes overall responsibility for that group. We suggest that the morphospecies reference image database is similarly managed by teams focused at the taxonomic grouping level. The appropriate taxonomic grouping will vary depending on variety represented by each phylogenetic level of the group, and expertise available. For example, Hexacorallia may have separate teams grouped at the Order level (e.g. Scleractinia, Actiniaria, Antipatharia), whereas Echinodermata may have separate teams grouped at Class level (e.g. Asteroidea, Echinoidea, etc.). Each team will consist of experts in taxonomy of the group plus ecologists engaged in field identification of organisms from imagery. We felt it was important to have both taxonomists and field ecologists working together, to ensure that the final database considers both taxonomic rigor and the practical use of the images. Each team will have a nominated lead, and leads will come together, as a steering committee, to ensure that a standard approach to data organisation and curation is achieved across the entire database.

We anticipate a two-stage process whereby an initial effort is made to collate and compile existing morphospecies reference image catalogues at a regional level using the new database structure described above. This would be followed by new and on-going submissions of data, including from those encountering new organisms not in the existing database, and from those with higher quality images of organisms already listed in the database (Fig. 2). We have committed to stage 1 of this process and morphospecies reference image databases held by all authors have been entered into this new database format and submitted to a central repository. Curation teams will now be established to bring these data submissions together into a single database that can be made available to end users.

**Figure 2:**
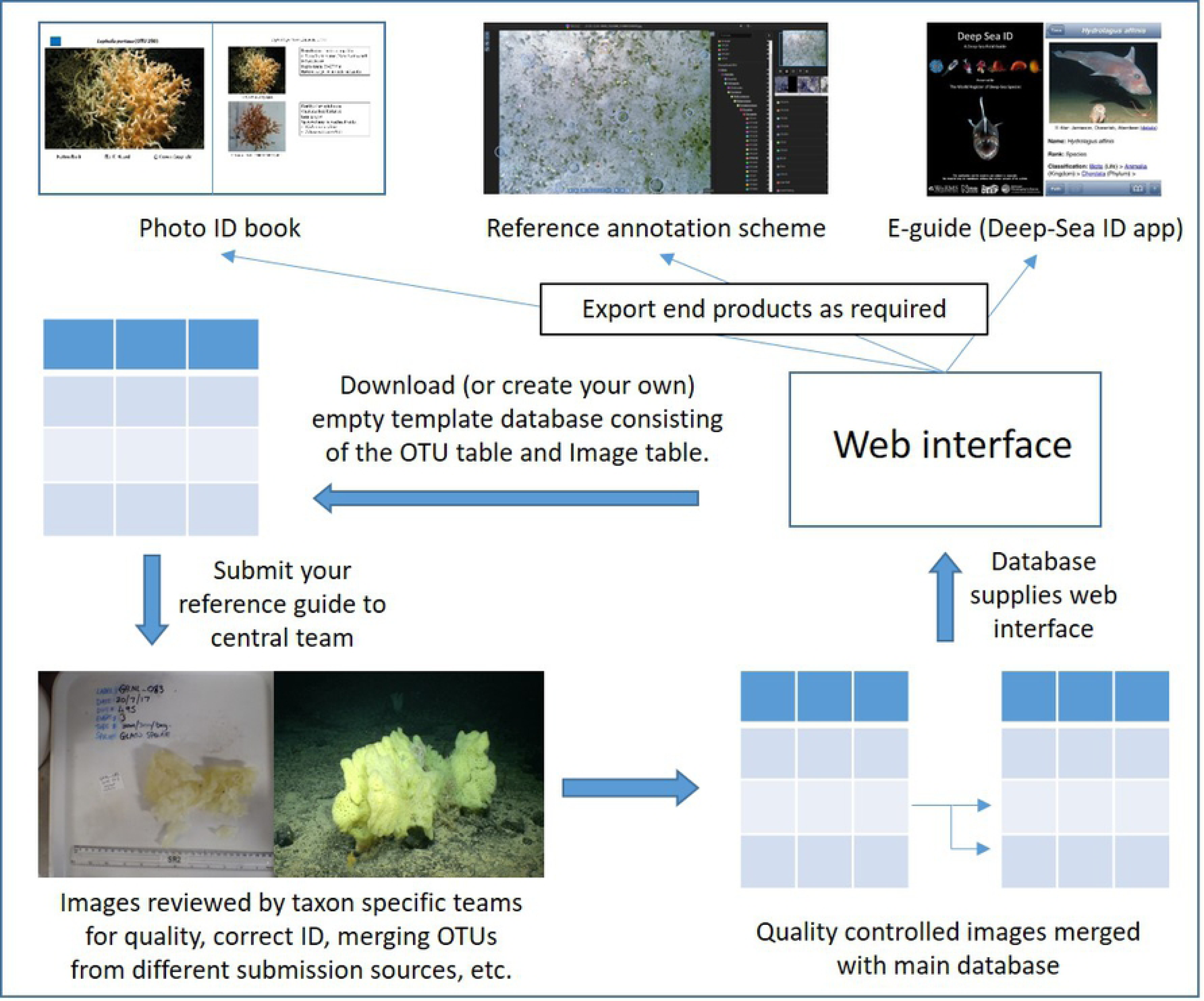
a conceptual model for how the developed framework will operate.

Stage 2 of this process will involve the effort of the global community and could potentially be a focus for the up-coming UN Decade of Ocean Science for Sustainable Development (2021-2030). This could be a very light-touch involvement, where end users simply submit images of new organisms not currently present in the database to the database for inclusion (Fig 2). Or it could be a more targeted and active involvement aimed at raising the quality of the data already in the database. For example, principal investigators of research cruises could actively help to move taxa from “identificationVerificationStatus” 1 to level 3 by targeted *in-situ* imaging and collection of organisms on an opportunistic basis.

Ultimately, it is not the database per-se that end users require, but the end products (photo-guides in book format, taxonomic tree for annotation software, etc.) that can be pulled from the database. This will require the development of a web interface that draws on the underlying database to produce multiple end use formats (Fig. 2). This aspect of the project represents the next stage of development and is anticipated to take place over the next two years.

## 4. Discussion

### 4.1. Immediate advances enabled by the development of a common reference standard

We have proposed a common structure for a database from which a morphospecies reference image catalogue can be built. Our initial development is focused on the North Atlantic deep-sea benthos as a case study. However, the structure developed is applicable to any marine region or habitat, and may also be used for terrestrial ecosystems. Individuals need only adopt the structure and populate the tables with their own data. The Standardised Marine Taxon Reference Image Database (SMarTaR-ID) will enable different researchers to bring their data together in a common morphospecies reference image catalogue at an appropriate time. Within the North Atlantic deep sea that time is now. The implementation of coherent monitoring programmes to assess biological biodiversity in marine waters are mandatory under the EU Marine Strategy Framework Directive (MSFD 2008/56/EC), and all European nations are required to monitor sites of community importance every six years. An image catalogue, such as the one herein proposed, will be a powerful instrument to support monitoring efforts, particularly in poorly surveyed regions. We have outlined a framework by which data can be brought together, curated, and new submissions managed going forward, which follows a successful model already applied by WoRMS.

We anticipate the introduction of a common reference standard for the deep sea to enhance significantly our understanding of megafaunal biodiversity by enabling multiple researchers to combine existing datasets to address long-standing ecological questions. This is particularly the case for hard substrate habitats that dominate features, such as seamounts, ridges, banks, abyssal hills, canyons, and areas of the continental slope, and for which image-based techniques remain the only effective means of survey. Past exploration of the deep-sea epibenthic megafauna generated many paradigms, but these were largely built on data obtained using trawls and sledges. Video and still image-based tools have facilitated quantitative sampling of previously inaccessible habitats; and the resulting new findings are challenging the prevailing view of deep-sea ecosystems [83]. However, these new datasets are often limited to individual features or feature types (e.g. seamounts: [84, 85], abyssal hills: [86] slopes: [66, 87, 88] canyons: [48, 64, 89, 90, 91]; ridges: [92], fracture zones: [85], and hydrothermal vents: [93]) and thus limit our ability to generalise findings. In their review of major outstanding questions in deep-sea biogeography [94] concluded, among other things, that an integrated biogeographic framework of hard-substrate areas of the deep sea was required to yield more realistic estimates of endemism/cosmopolitanism. It has been repeatedly argued that concerted efforts to link existing independent data streams together to examine long-standing questions of deep-sea diversity are very much needed in order to move the field forward [94, 95]. The proposed database will facilitate these advances.

We anticipate that this common reference standard will provide an invaluable tool for environmental managers, industry and wider stakeholders. For environmental managers, it will, for example, enable the development of clearer descriptions and definitions of habitats of conservation concern. For example, deep-sea sponge aggregations potentially qualify as Vulnerable Marine Ecosystems (VME) under the United Nations General Assembly (UNGA) Resolution 61/105. They are also classed as a threatened and declining ecosystem under Annex V of the Oslo-Paris (OSPAR) Convention for the Protection of the Marine Environment of the North East Atlantic. However, comprehensive descriptions of deep-sea sponge aggregations, and specifically the component taxa that compose different types of aggregation, are lacking. In addition, basin-wide data on the distribution of sponge VME indicator taxa are only available for those species / genera whose appearance both *in-situ* and *ex-situ* are similar (e.g. Geodia, Hyalonema, Pheronema). For many sponge species, the lack of taxonomic resolution possible when identifying sponges from image data hinders progress in management and conservation of these taxa by limiting our ability to 1) effectively describe sponge VME composition and diversity, and 2) pool data to determine basin-wide distributions. A common morphospecies reference image catalogue will provide a standard reference to use in VME descriptions in the absence of confirmed taxonomic identification of species from physical samples. It will also facilitate the production of basin-wide models of the distribution of habitat forming sponge taxa to support spatial management decisions [96].

For industry, implementation of a standard approach to referencing morphospecies between industry and regulators will facilitate a much more effective impact assessment associated with licensing and consent processes, as well as subsequent monitoring approaches. Often in industry, a range of sub-contractors are used for routine survey and monitoring work by the various industry bodies. Therefore, morphotypes are produced per project with no consistency between sub-contractor or between years in long-term monitoring as data are rarely shared. This standardisation would increase industry and regulatory comparison across applications and across industries to facilitate cumulative impact assessments, thus allowing better understanding of impact at feature and site levels, as required in nature conservation legislation. For industry, this could also decrease levels of risk associated with the assessments as well as decreased analysis time and costs for survey data, and would be a particularly powerful tool if industry could include their own data in the database and play an active role in providing images and survey data.

The need for a standard approach in industry was recently highlighted by the development of the deep-sea mining industry in the Clarion Clipperton Zone (CCZ) of the central Pacific. Here, baseline data collection is taking place, commonly including seabed imaging-based assessments of megafauna [29, 47, 65, 97, 98]. Without a consistent morphospecies reference image catalogue it is difficult to compare studies and generate regional syntheses. This greatly hampers conservation and management efforts, which commonly rely on information on biodiversity, species ranges and behaviour – ecological properties that are difficult to assess without good quality and consistent identifications. Recent work to document megafaunal diversity will help (e.g. 47, 65), but widely adopted and regularly updated catalogues will be vital for improving scientific understanding and effective environmental management.

A common morphospecies reference image catalogue may also serve as a tool to support the identification of taxa from fisheries bycatch by fisheries observers (e.g. [99]. While our proposed database focuses on *in-situ* images of taxa, we advocate, and have provided for within the proposed database structure, the collection of *ex-situ* images of taxa. There are a number of existing image guides designed for use by fisheries observers that provide *ex-situ* images of VME indicator taxa (Table 1). This database could supplement existing guides by providing additional imagery. Interestingly, it may also provide a link between *in-situ* and *ex-situ* taxon identification, which may ultimately allow fisheries bycatch data to be pooled with *in-situ* image data, again broadening our understanding of species distributions (e.g. [100]).

Finally, the simple act of combining multiple existing morphospecies reference image catalogues will advance the overall quality of current identifications. Different research groups have images of different species for which the “identificationVerificationStatus” level is 3 (the highest level, confirmed by physical specimen). By bringing these reference image sets together, we will collectively have more species that can be identified by reference to images in which we have the highest level of confidence of the animal’s identification.

### 4.2. Future advances enabled by the development of a common reference standard

The development of a common reference standard has the potential to advance significantly the field of offshore and deep-sea marine ecology. The ability to pool datasets across time and space will allow us to address a greater range of questions about the offshore and deep-sea benthic ecosystem than is currently possible. Critically, it will enable us to raise standards of identification from image data through the development of training materials and quality control measures. Efforts to develop such tools for shallow water have been undertaken by the UK’s National Marine Biological Association Quality Control scheme (NMBAQC). This programme is steered by a range of academic and governmental organisations, and provides guidance on best practice, as well as identification guides, taxonomic workshops, training exercises and quality control ring tests.

There will remain some potential shortcomings on the use of such catalogues related to uncertainties in species identification due to the method of image collection and scale. The ability to zoom-in on specific features of species with ROV cameras means ROVs may provide better imagery for identification than AUVs or drop-down cameras, particularly in cases where species look remarkably similar and occupy overlapping environmental niches. For example, the octocorals *Acanthogorgia armata* and *Acanthogorgia hirsuta* can only be distinguished if close up images of the polyps are taken, otherwise identifications have to be left at genus level. Nevertheless, the development of a common reference standard will expose these limitations to a wider audience, and help develop agreed international guidance around the taxonomic levels to which it is appropriate to identify when interpreting image data.

In the longer term, regional field keys are required for use in survey and monitoring of the deep-sea ecosystem. The construction of tools that allow others to identify taxa reliably and consistently in the field is perhaps one of the most underappreciated roles for taxonomists. It is also one of the most challenging roles as taxonomists are often not engaged in field identification, and therefore a gap exists between the generator and end user of a key [101]. The starting point for the development of any key is a standard reference against which to compare new observations. In traditional taxonomy, this is the type specimen, a physical specimen from which a species is described, that is subsequently archived in a museum. The development of dichotomous or polytomous keys is then achieved by measuring the variability in observable characteristics within examples of a taxon and between taxa, then selecting characteristics that best discriminate between taxa for a given region / group. These characteristics are then organised into pathways of character state choices (steps) that lead to identifications.

In order to move forward with the development of much-needed field keys to deep-water taxa, we must first develop an appropriate standard reference against which to assess new observations. This reference point remains the primary type specimen. Our proposed database will establish an image ‘similitype’ (or a series of images that contribute to the similitype) to accompany a physical specimen that has been identified with reference to the primary type of a species, or through matching DNA sequences to other specimens identified with reference to the primary type of a species, and thus link traditional taxonomy to field identification. This approach will provide a much-needed strategy to advance the taxonomic description of species based on multisource information collected by both ecologists and taxonomists [102, 103]. If researchers use and contribute to this common reference standard, a library of images with examples of each taxon will be built up over time. This library of image examples can then be used to understand both the observable characteristics within a species or higher taxonomic level grouping, and the variability in these characteristics in image-based data. Where possible it is desirable for these characteristics to be those used in traditional taxonomic keys. However, this will not be possible for all groups to all levels of the taxonomic hierarchy. For example, while it is possible to use traditional taxonomic features to determine the order of some coral taxa from image data, it is not possible to do this for anemone taxa, which rely on internal characteristics for positive identification. It is likely that novel characteristics, combinations of characteristics, as well as the use of circumstantial information (e.g. environmental characteristics), will be required to enable reliable and consistent field identification of organisms.

Multi-access keys (also known as matrix based or free-access keys) may be more appropriate than dichotomous or polytomous keys (also known as single-access keys) for use with image data as they, by their nature, have multiple access points [101, 104]. Single-access keys place a logical order on the use of characteristics, with each step in the decision tree taking the user along a predefined pathway that progressively narrows the number of possibilities for the identification of the animal. If a characteristic is not visible at any step along this pathway, the choice required by the user is unanswerable and further progression is not possible. Views of organisms in *in-situ* image data can be highly variable, and it is likely that in any one image only some features will be visible. This may limit the utility of single-access keys with image data. Multi-access keys enable the user to determine the sequence of choices where the user can select from the list of characteristics offered in order to arrive at an identification. In the context of image data, this would allow the user to employ all visible characteristics (and potentially environmental information) to arrive at an identification. Multi-access keys are more suitable for computer-aided identification tools [101, 104]. This also makes them a promising tool to use with image data where analysis is computer based.

If we are to move forward with the application of AI and CV to the identification of taxa, we must have a common reference standard. Our proposed database aims to meet this need through future development that will enable the database to interface with image annotation software, such as Squidle [105] and BIIGLE 2.0 [106; 107]. These annotation softwares enable users to mark the x,y position of organisms within an image and attribute this point / polygon with a taxon identification. This process of image annotation is the means by which ecologists extract semantic data from an image in order to then apply numerical and statistical analysis to these data and answer ecological questions. This annotated dataset is also the base data needed in the development of AI and CV algorithms. These algorithms require large numbers of images to “learn” the features that distinguish the different OTUs to which they have been exposed, and which of these features are characteristic of each OTU [108, 109]. If researchers are able to use a common reference standard, thus extracting the same information from an image regardless of who is annotating it, then collated datasets from various origins could reach the size needed to train and test CV algorithms (acknowledging challenges of observer bias). Their use within the field of deep-sea benthic ecology, will then increase exponentially through accumulation of data, skill and experience. This can only serve to facilitate the development of CV and bring us closer to automation of image annotation and data extraction.

Ultimately, standardisation of tools and methods is central to long-term monitoring and assessments of ocean health. Woodall et al. [110] recognised this and produced GOSSIP (General Ocean Survey and Sampling Iterative Protocol), which outlines a framework of 20 biological, chemical, physical, and socioeconomic parameters that allow marine scientists to generate comparable data on the function, health and resilience of the ocean. There are several international efforts underway to try and harmonise ocean observing in the areas of biology and ecology, including the efforts of the Group on Earth Observation – Biodiversity Observing Network (GEO-BON) and the Global Ocean Observing System (GOOS) panel on biology and ecosystem – Essential Ocean Variables (EOVs). These efforts are also being informed by international efforts, such as the Deep Ocean Observing Strategy (http://www.deepoceanobserving.org/), which is adding deep ocean context to GOOS EOV specifications. There are more than a dozen regional alliances internationally, which are implementing the GOOS vision with international coordination by the IOC. Together these organisations are forming a means for efforts from individual observers, as well as local to international bodies, to join together to realise the power of ‘big data’ in observing and understanding change. National-level data management and communications groups affiliated with GOOS are now working to include tools, such as automated image classification, into their information technology systems. The common reference image standard described above will therefore contribute to global efforts under GOOS.

## 5. Conclusions

We have developed a database structure to facilitate the standardisation of morphospecies image catalogues between individuals, research groups, and nations. We have also proposed a framework for coordination of international efforts to develop reference guides for the identification of deep-sea species from images. We have highlighted the potential gains to be made through the use of this database structure by the deep-sea community in: increasing the quality and quantity of data available to researchers, improvement of overall understanding of the deep-sea ecosystem, more effective management and monitoring by statutory bodies and industry alike, and realising the potential benefits of emerging AI and CV approaches. To make these gains it is critical there is now uptake of this database structure by the community, and additional funding is found to contribute to stage two development.

## Acknowledgements

KH, NP, RR, NF was supported by the NERC funded DeepLinks project NE/K011855/1. The workshop was funded by the Deep Sea Biology Society’s Lounsbery Workshop Award. ALA and CLM are supported by Grant Number SFI/15/IA/3100 to ALA from Science Foundation Ireland and the Marine Institute under the Investigators Programme co-funded under the European Regional Development Fund 2014-2020. AB-H was supported by the Oceanic Observatory of Madeira project (M1420-01-0145-FEDER-000001-Observatório Oceânico da Madeira-OOM) co-financed by the Madeira Regional Operational Programme (Madeira 14-20) under the Portugal 2020 strategy through the European Regional Development Fund, and the Portuguese Foundation for Science and Technology (FCT, Portugal), through the strategic project UID/MAR/04292/2013 granted to MARE. JV is supported by Oil and Gas UK and the ATLAS project funded by the European Commission’s H2020 Scheme through Grant Agreement 678760. HAR was supported by the CeNCOOS Partnership: Ocean Information for Decision Makers (award number NA16NOS0120021). DOBJ was supported by the UK Natural Environment Research Council National Capability funding: “Climate Linked Atlantic Section Science” (CLASS), grant number NE/R015953/1. DW was supported by NOAA Deep Sea Coral Research and Technology Program. LW and PS were supported by the Garfield Weston Foundation. TM was supported by Program Investigador FCT (IF/01194/2013), IFCT Exploratory Project (IF/01194/2013/CP1199/CT0002) from the Fundação para a Ciência e Tecnologia (POPH and QREN), PO2020 MapGes (Acores-01-0145-FEDER-000056), and H2020 ATLAS (grant agreement no. 678760). RV was funded by the Fundação para a Ciência e a Tecnologia (FCT/SFRH/BD/84030/2012), with additional support provided by Cefas through the Science Futures programme. JRX research is funded by the H2020 EU Framework Programme for Research and Innovation through the SponGES project (grant agreement No. 679849) and partially supported by the Strategic Funding UID/Multi/04423/2019 through national funds provided by the Foundation for Science and Technology (FCT) and the European Regional Development Fund (ERDF), in the framework of the programme PT2020.

## References

1. Ruppé CV, Barstad JF, editors. International handbook of underwater archaeology. Berlin: Springer Science & Business Media; 2013.

2. Boutan L. La photographie sous-marine. Arch Zool Exp. 1893;3: 281–324.

3. Cousteau JY. The living sea. London: H. Hamilton; 1963.

4. Beebe W. Half mile down. Duell, Sloan and Pearce; 1951.

5. Ewing M, Vine A, Worzel JL. Photography of the ocean bottom. JOSA. 1946 Jun 1;36(6):307–21.

6. Ewing M, Worzel JL, Vine AC. Early development of ocean-bottom photography at Woods Hole Oceanographic Institution and Lamont Geological Observatory. The John Hopkins Oceanographic Studies. 1967.

7. Schenck HJ, Kendall H. Underwater photography. Maryland: Cornell Maritime Press; 1954.

8. Thorndike EM. Deep-sea cameras of the Lamont Observatory. Deep Sea Research (1953). 1958 Jan 1;5(2-4):234–7.

9. Fell HB. Biological applications of sea-floor photography. In: Hersey JB, editor. Deep-sea photography. Baltimore: John Hopkins Press. 1967. pp 207–221.

10. Vevers HG. Photography of the sea floor. J Mar Biol Assoc UK 1951;30: 101–111.

11. Clark HES. Fauna of the Ross Sea Part 3: Asteroidea. Mem N Z Oceanogr Inst 1963;21: 1–84.

12. Marshall NB, Bourne DW. A photographic survey of benthic fishes in the Red Sea and Gulf of Eden, with observations on their population density, diversity and habitats. Bull Mus Comp Zool 1964;132: 225–244.

13. Hersey JB. Deep-sea photography. Baltimore: John Hopkins Press; 1967.

14. Heezen BC, Hollister CD. The face of the deep. London: Oxford University Press; 1971.

15. Grassle JP, Sanders RR, Hessler GT, Rowe GT, McLellan T. Pattern and zonation: a study of the bathyal megafauna using the research submersible Alvin. Deep Sea Res I 1975;22: 457–481.

16. Rice AL, Aldred G, Darlington E, Wild RA. The quantitative estimation of the deep-sea megabenthos: a new approach to an old problem. Oceanol Acta 1982;5: 63–72.

17. Rowe GT, Sibuet M, Vangriesheim A. Domains of occupation of abyssal scavengers inferred from baited cameras and traps on the Demerara Abyssal Plain. Deep Sea Res Part I 1986;33: 501–522.

18. Smith KL, Kaufmann RS, Wakefield WW. Mobile megafaunal activity monitored with a time-lapse camera in the abyssal North Pacific. Deep Sea Res I 1993;40: 2307–2324.

19. Thurston MH, Bett BJ, Rice AL, Jackson PAB. Variations in the invertebrate abyssal megafauna in the North Atlantic Ocean. Deep Sea Res I 1994;41: 1321–1348.

20. Howell KL, Davies J, Hughes DJ, Narayanaswamy BE. Strategic Environmental Assessment / Special Area for Conservation Photographic Analysis Report. London: Department of Trade and Industry; 2007.

21. Durden JM, Schoening T, Althaus F, Friedman A, Garcia R, Glover AG, Greinert J, Jacobsen Stout N, Jones DOB, Jordt A, Kaeli JW, Koser K, Kuhnz LA, Lindsay D, Morris KJ, Nattkemper TW, Osterloff J, Ruhl HA, Singh H, Tran M, Bett BJ. Perspectives in visual imaging for marine biology and ecology: from acquisition to understanding. Oceanogr Mar Biol Annu Rev 2016;54: 1–72.

22. Taylor J, Krumpen T, Soltwedel T, Gutt J, Bergmann M. Dynamic benthic megafaunal communities: Assessing temporal variations in structure, composition and diversity at the Arctic deep-sea observatory HAUSGARTEN between 2004 and 2015. Deep-Sea Res I 2017;122: 81–94.

23. Taylor J, Krumpen T, Soltwedel T, Gutt J, Bergmann M. Regional- and local-scale variations in benthic megafaunal composition at the Arctic deep-sea observatory HAUSGARTEN. Deep Sea Res I 2016;108: 58–72.

24. Howell KL, Davies JS, Narayanaswamy BE. Identifying deep-sea megafaunal epibenthic assemblages for use in habitat mapping and marine protected area network design. J Mar Biol Assoc UK 2010a;90: 33–68.

25. Huvenne VAI, Bett BJ, Masson DG, Le Bas TP, Wheeler AJ. Effectiveness of a deep-sea cold-water coral Marine Protected Area, following eight years of fisheries closure. Biol Conserv 2016;200: 60–69.

26. Escartin J, Barreyre T, Cannat M, Garcia R, Gracias N, Deschamps A, Salocchi A, Sarradin PM, Ballu V. Hydrothermal activity along the slow-spreading Lucky Strike ridge segment (Mid-Atlantic Ridge): Distribution, heatflux, and geological controls. Earth Planet Sci Lett 2015;431: 173–185.

27. Hirai J, Jones DOB. The temporal and spatial distribution of krill (Meganyctiphanes norvegica) at the deep seabed of the Faroe–Shetland Channel, UK: A potential mechanism for rapid carbon flux to deep sea communities. Mar Biol Res 2011;8: 48–60.

28. Olu K, Lance S, Sibuet M, Henry P, Fiala-Médioni A, Dinet A. Cold seep communities as indicators of fluid expulsion patterns through mud volcanoes seaward of the Barbados accretionary prism. Deep Sea Res I 1997;44: 811–819.

29. Simon-Lledó E, Bett BJ, Huvenne VAI, Schoening T, Benoist NMA, Jeffreys RM, Durden JM, Jones DOB. Megafaunal variation in the abyssal landscape of the Clarion Clipperton Zone. Prog Oceanogr 2019a;170: 119–133.

30. Laurenson C, Hudson IR, Jones DOB, Preide IM. Deep water observations of *Lophius piscatorius* in the north-eastern Atlantic Ocean by means of a Remotely Operated Vehicle. Fish Biol 2004;65: 947–960.

31. Jones DOB, Bett BJ, Tyler PA. Megabenthic ecology of the Faroe-Shetland Channel: a photographic study. Deep Sea Res I. 2007;54: 1111–1128.

32. Durden JM, Bett BJ, Ruhl HA. The hemisessile lifestyle and feeding strategies of *Iosactis vagabunda* (Actiniaria, Iosactiidae), a dominant megafaunal species of the Porcupine Abyssal Plain. Deep Sea Res I. 2015a;102: 72–77.

33. Bullimore RD, Foster NL, Howell KL. Coral-characterized benthic assemblages of the deep Northeast Atlantic: defining “Coral Gardens” to support future habitat mapping efforts. ICES J Mar Sci. 2013;70: 511–522.

34. Morato TM, Pham CK, Pinto C, Golding N, Ardron JA, Durán Muñoz P, Neat F. A multi criteria assessment method for identifying Vulnerable Marine Ecosystems in the North-East Atlantic. Front Mar Sci. 2018;5: 460.

35. Pham CK, Diogo H, Menezes G, Porteiro F, Braga-Henriques A, Vandeperre F, Morato T. Deep-water longline fishing has reduced impact on Vulnerable Marine Ecosystems. Sci Rep. 2014a;4:4837.

36. Buhl-Mortensen P. Coral reefs in the Southern Barents Sea: habitat description and the effects of bottom fishing. Mar Biol Res. 2017;13: 1027–1040.

37. Pham CK, Ramirez-Llodra E, Alt CHS, Amaro T, Bergmann M, Canals M, Company JB, Davies J, Duineveld G, Galgani F, Howell KL, Huvenne VAI, Isidro E, Jones DOB, Lastras G, Morato T, Gomes-Pereira JN, Purser A, Stewart H, Tojeira I, Tubau X, Van Rooij D, Tyler PA. Marine litter distribution and abundance in European Seas, from the shelf to deep basins. PLOS ONE. 2014b;9: e95839.

38. Buhl-Mortensen P, Buhl-Mortensen L. Impacts of Bottom Trawling and Litter on the Seabed in Norwegian Waters. Front Mar Sci. 2018;5:42 doi:10.3389/fmars.2018.00042

39. Billett DSM, Bett BJ, Reid WDK, Boorman B, Priede IG. Long-term change in the abyssal NE Atlantic: the ‘Amperima Event’ revisited. Deep Sea Res II 2010;57: 1406–1417.

40. Morris KJ, Bett BJ, Durden JM, Huvenne VAI, Milligan R, Jones DOB, McPhail S, Robert K, Bailey DM, Ruhl HA. A new method for ecological surveying of the abyss using autonomous underwater vehicle photography. Limnol Oceanogr Methods. 2014;12: 795–809.

41. Edgar GJ. Australian marine life: the plants and animals of temperate waters. Sydney: Reed New Holland; 2008.

42. Wood C. Sea anemones and corals of Britain and Ireland. Plymouth: Wild Nature Press; 2013.

43. Jacobsen Stout N, Kuhnz L, Lundsten L, Schlining B, Schlining K, von Thun S. The Deep-Sea Guide (DSG). Monterey Bay Aquarium Research Institute (MBARI). 2015. Available from: http://dsg.mbari.org/dsg/home

44. Braga-Henriques A, Pereira JN, Tempera F, Porteiro FM, Pham C, Morato T, Santos RS. Cold-water coral communities on Condor Seamount: initial interpretations. In: Giacomello E, Menezes G (eds) CONDOR observatory for long-term study and monitoring of azorean seamount ecosystems. Final Project Report, Arquivos do DOP, Série Estudos 1/2012, Horta. 2011. pp 105–114.

45. Braga-Henriques A, Carreiro-Silva M, Tempera F, Porteiro FM, Jakobsen K, Jakobsen J, Albuquerque M, Santos RS. Carrying behavior in the deep-sea crab *Paromola cuvieri* (Northeast Atlantic). Mar Biodiv. 2012;42: 37–46

46. Narayanaswamy BE, Hughes DJ, Howell KL, Davies J, Jacobs C. First observations of megafaunal communities inhabiting George Bligh Bank, northeast Atlantic. Deep Sea Res II. 2013;92: 79–86.

47. Amon DJ, Ziegler A, Kremenetskaia A, Mah C, Mooi R, O’Hara T, Pawson D, Roux M, Smith C. Megafauna of the UKSRL exploration contract area and eastern Clarion-Clipperton Zone in the Pacific Ocean: Echinodermata. Biodivers Data J. 2017a;5: e11794.

48. van den Beld IMJ, Bourillet JF, Arnaud-Haond S, de Chambure L, Davies JS, Guillaumont B, Olu K, Menot L. Cold-water coral habitats in submarine canyons of the Bay of Biscay. Front Mar Sci. 2017;4: 10.3389/fmars.2017.00118

49. Alt CHS, Kremenetskaia A, Gebruk AV, Gooday AJ, Jones DOB. Bathyal benthic megafauna from the Mid-Atlantic Ridge in the region of the Charlie-Gibbs fracture zone based on remotely operated vehicle observations. Deep Sea Res I. 2019;145: 1–12.

50. Hawkes N, Korabik M, Beazley L, Rapp HT, Xavier JR, Kenchington E. Glass sponge grounds on the Scotian Shelf and their associated biodiversity. Mar Ecol Prog Ser. 2019;614: 91–109.

51. Culverhouse PF, Williams R, Reguera B, Herry V, Gonzalez-Gils. Do experts make mistakes? A comparison of human and machine identification of dinoflagellates. Mar Ecol Prog Ser. 2003;247: 17–25.

52. MacLeod N, Benfield M, Culverhouse P. Time to automate identification. Nature. 2010;467: 154–155.

53. Schoening T, Bergmann M, Ontrup J, Taylor J, Dannheim J, Gutt J, Nattkemper TW. Semi-automated image analysis for the assessment of megafaunal densities at the Arctic deep-sea observatory HAUSGARTEN. PLOS ONE. 2012;7: e38179.

54. Wynn RB, Huvenne VAI, Le Bas TP, Murton BJ, Connelly DP, Bett BJ, Ruhl HA, Morris KJ, Peakall J, Parsons DR, Sumner EJ, Darby SE, Dorrell RM, Hunt JE. Autonomous Underwater Vehicles (AUVs): their past, present and future contributions to the advancement of marine geoscience. Mar Geol. 2014;352: 451–468.

55. Jones DOB, Gates AR, Huvenne VAI, Phillips AB, Bett BJ. Autonomous marine environmental monitoring: Application in decommissioned oil fields. Sci Total Environ. 2019;668: 835–853.

56. Piechaud N, Hunt C, Culverhouse PF, Foster NL, Howell KL. Automated identification of benthic epifauna with computer vision. Mar Ecol Prog Ser. 2019;615: 15–30.

57. . Edgington DR, Cline DE, Davis D, Kerkez I, Mariette J. Detecting, tracking and classifying animals in underwater video. Proc Oceans IEEE. 2006.

58. Beijbom O, Edmunds PJ, Roelfsema C, Smith J, Kline DI, Neal BP, Dunlap MJ, Moriarty V, Fan T-Y, Tan C-J. Towards automated annotation of benthic survey images: Variability of human experts and operational modes of automation. PLOS ONE. 2015;10:e0130312.

59. Schoening T, Durden J, Preuss I, Albu AB, Purser A, De Smet B, Dominguez-Carrió C, Yesson C, de Jonge D, Lindsay D. Report on the marine imaging workshop 2017. Res Ideas Outcomes. 2017;3:e13820.

60. Favret C, Sieracki JM. Machine vision automated species identification scaled towards production levels. Syst Entomol. 2016;41: 133–143.

61. Langenkämper D, Nattkemper TW. COATL – A learning architecture for online real-time detection and classification assistance for environmental data. IEEE Int Conf Pattern Recognit, IEEE, 2017a. pp 597–602.

62. Howell KL, Davies JS. Deep-sea species image catalogue, On-line version 2. 2016. Available from: https://deepseacruorg/2016/12/16/deep-sea-species-image-catalogue/

63. Jones DOB, Gates AR. Deep-sea life of Scotland and Norway. UK: Ophiura; 2010.

64. Robert K, Jones DOB, Tyler PA, Van Rooji D, Huvenne VAI. Finding the hotspots within a biodiversity hotspot: fine-scale biological predictions within a submarine canyon using high-resolution acoustic mapping techniques. Mar Ecol. 2014;36: 1256–1276.

65. Amon DJ, Ziegler AF, Drazen JC, Grischenko AV, Leitner AB, Lindsay DJ, Voight JR, Wicksten MK, Young CM, Smith CR. Megafauna of the UKSRL exploration contract area and eastern Clarion-Clipperton Zone in the Pacific Ocean: Annelida, Arthropoda, Bryozoa, Chordata, Ctenophora, Mollusca. Biodivers Data J. 2017b;5: e14598–e14598

66. Stefanoudis P, Smith S, Schneider C, Wagner D, Goodbody-Gringley G, Xavier J, Rivers M, Woodall L, Rogers A. Deep Reef Benthos of Bermuda: Field Identification Guide. Figshare Book. 2018. Available from: https://doi.org/10.6084/m9.figshare.7333838.v1

67. Althaus F, Hill N, Ferrari R, Edwards L, Przeslawski R, Schönberg CH, Stuart-Smith R, Barrett N, Edgar G, Colquhoun J. A standardised vocabulary for identifying benthic biota and substrata from underwater imagery: the CATAMI classification scheme. PLOS ONE. 2015;10:e0141039

68. Wieczorek J, Bloom D, Guralnick R, Blum S, Doring M, Giovanni R, Robertson T, Vieglais D. Darwin Core: an evolving community-developed biodiversity data standard. PLOS ONE. 2012;7: e2971569.

69. Encyclopedia of Life. Available from http://www.eol.org

70. GBIF.org. GBIF Home Page. 2018. Available from https://www.gbif.org

71. OBIS. Ocean Biogeographic Information System. Intergovernmental Oceanographic Commission of UNESCO. 2018. Available from www.iobis.org.

72. WoRMS Editorial Board. World Register of Marine Species. 2018. Available from http://www.marinespecies.org.

73. Vandepitte L, Vanhoorne B, Decock W, Vranken S, Lanssens T, Dekeyzer S, Verfaille K, Horton T, Kroh A, Hernandez F, Mees J. A decade of the World Register of Marine Species – General insights and experiences from the Data Management Team: Where are we, what have we learned and how can we continue? PLOS ONE 2018;13: e0194599

74. Horton T, Gofas S, Kroh A, Poore GCB, Read G, Rosenberg G, Stöhr S, Bailly N, Boury-Esnault N, Brandão SN, Costello MJ, Decock W, Dekeyzer N, Hernandez F, Mees J, Paulay G, Vandepitte L, Vanhoorne B, Vranken S. Improving nomenclatural consistency: a decade of experience in the World Register of Marine Species. Eur J Taxon. 2017;389: 1–24.

75. Glover AG, Higgs ND, Horton T, Porrer A. Deep Sea ID v.1.2 A Field Guide to the Marine Life of the Deep Sea 2015. Available from http://www.marinespecies.org/deepsea

76. Claus S, De Hauwere N, Vanhoorne B, Souza Dias F, Oset García P, Schepers L, Hernandez F, Mees J. MarineRegions.org. 2018. Available from http://www.marineregions.org

77. Howell KL, Billett DSM, Tyler PA. Depth-related distribution and abundance of seastars (Echinodermata : Asteroidea) in the Porcupine Seabight and Porcupine Abyssal Plain, NE Atlantic. Deep Sea Res I. 2002;49: 1901–1920.

78. Greene HG, Yoklavich MM, Starr RM, O’Connell VM, Wakefield WW, Sullivan DE, McRea JE, Cailliet GM. A classification scheme for deep seafloor habitats. Oceanol Acta 1999;22: 663–678.

79. Davies CE, Moss D. EUNIS Habitat Classification. Final Report to the European Topic Centre on Nature Conservation, European Environment Agency, Copenhagen; 1998.

80. Davies CE, Moss D, Hill MO. EUNIS Habitat Classification Revised 2004. Report to the European Topic Centre on Nature Protection and Biodiversity, European Environment Agency, Copenhagen; 2004.

81. Folk RL. The distinction between grain size and mineral composition in sedimentary rock nomenclature. J Geol. 1954;62: 344–359.

82. Wentworth CK. A scale of grade and class terms for clastic sediments. J Geol. 1922;30: 377–392.

83. Danovaro R, Snelgrove PV, Tyler P. Challenging the paradigms of deep-sea ecology. Trends Ecol Evol. 2014;29: 465–475.

84. Howell KL, Mowles SL, Foggo A. Mounting evidence: near-slope seamounts are faunally indistinct from an adjacent bank. Mar Ecol – Evol Persp. 2010b;31: 52–62.

85. Victorero L, Robert K, Robinson LF, Taylor ML, Huvenne VAI. Species replacement dominates megabenthos beta diversity in a remote seamount setting. Sci Rep. 2018;8: 4152.

86. Durden JM, Bett BJ, Jones DOB, Huvenne VAI, Ruhl HA. Abyssal hills a hidden source of increased habitat heterogeneity, benthic megafaunal biomass and diversity in the deep sea. Prog Oceanogr. 2015b;137: 209–218.

87. Buhl-Mortensen L, Buhl-Mortensen P, Dolan MFJ, Dannheim J, Bellec V, Holte B. Habitat complexity and bottom fauna composition at different scales on the continental shelf and slope of northern Norway. Hydrobiologia. 2012;685 :191–219.

88. Fonseca P, Abrantes F, Aguilar R, Campos A, Cunha M, Ferreira D, Fonseca TP, Garcia S, Henriques V, Machado M, Mecho A, Relvas P, Rodrigues CF, Salgueiro E, Vieira R, Weetman A, Castro M. A deep-water crinoid *Leptometra celtica* bed off the Portuguese south coast. Mar Biodivers. 2014;44: 223–228.

89. Huvenne VAI, Tyler PA, Masson DG, Fisher EH, Hauton CH, Hühnerbach V, Le Bas TP, Wolff GA. A picture on the wall: Innovative mapping reveals cold-water coral refuge on submarine canyon. PLOS ONE. 2011;6: e28755.

90. Johnson MP, White M, Wilson A, Würzberg L, Schwabe E, Folch H, Allcock AL. A vertical wall dominated by *Acesta excavata* and *Neopycnodonte zibrowii*, part of an undersampled group of deep-sea habitats. PLOS ONE. 2013;8: e79917

91. Davies JS, Howell KL, Stewart HA, Guinan J, Golding N. Defining biological assemblages (biotopes) of conservation interest in the submarine canyons of the South West Approaches (offshore United Kingdom) for use in marine habitat mapping. Deep Sea Res II. 2014;104: 208–229.

92. Bell JB, Alt CHS, Jones DOB. Benthic megafauna on steep slopes at the Northern Mid-Atlantic Ridge. Mar Ecol. 2016;37: 1290–1302.

93. Marsh L, Copley JT, Huvenne VAI, Tyler PA and the Isis ROV Facility. Getting the bigger picture: Using precision Remotely Operated Vehicle (ROV) videography to acquire high-definition mosaic images of newly discovered hydrothermal vents in the Southern Ocean. Deep Sea Res II. 2013;92: 124–135.

94. McClain CR, Hardy SM. The dynamics of biogeographic ranges in the deep sea. Proc R Soc Lond [Biol]. 2010;277: 3533–3546.

95. McClain CR, Schlacher TA. On some hypotheses of diversity of animal life at great depths on the sea floor. Mar Ecol. 2015;36: 849–872.

96. Howell KL, Piechaud N, Downie AL, Kenny A. The distribution of deep-sea sponge aggregations in the North Atlantic and implications for their effective spatial management. Deep Sea Res I. 2016;115: 309–320.

97. Vanreusel A, Hilario A, Ribeiro PA, Menot L, Arbizu PM. Threatened by mining, polymetallic nodules are required to preserve abyssal epifauna. Sci Rep. 2016;6: 26808

98. Simon-Lledó E, Bett BJ, Huvenne VAI, Schoening T, Benoist NMA, Jones DOB. Ecology of a polymetallic nodule occurrence gradient: Implications for deep-sea mining. Limnol Oceanogr 2019b.

99. CCAMLR. VME Taxa Classification Guide. Commission for the Conservation of Antarctic Marine Living Resources, Hobart, Tasmania, 2009; 4pp

100. Vieira RP, Cunha MR. *In situ* observation of chimaerid species in the Gorringe Bank: new distribution records for the north-east Atlantic Ocean. J Fish Biol. 2014;85: 927–932.

101. Walter DE, Winterton S. Keys and the crisis in taxonomy: extinction or reinvention? Annu Rev Entomol. 2007;52: 193–208.

102. Grandcolas P. Loosing the connection between the observation and the specimen: a by-product of the digital era or a trend inherited from general biology? Bionomina. 2017;12: 57–62.

103. Thomson SA, Pyle RL, Ahyong ST, Alonso-Zarazaga M, Ammirati J, Araya JF, Ascher JS, Audisio TL, Azevedo-Santos VM, Bailly N, Baker WJ. Taxonomy based on science is necessary for global conservation. PLOS Biol. 2018;16: e2005075

104. Hagedorn G, Rambold G, Martellos S. Types of identification keys. In Nimis PL, Vignes Lebbe R, editors. Tools for identifying biodiversity: progress and problems. Proc Int Cong Paris, Edizioni Università di Trieste 2012; pp 59–64.

105. Williams S, Friedman A. SQUIDLE+ 2018. Available from: http://squidle.acfr.usyd.edu.au.

106. Ontrup J, Ehnert N, Bergmann M, Nattkemper TW. BIIGLE – Web 2.0 enabled labelling and exploring of images from the Arctic deep-sea observatory HAUSGARTEN. In OCEANS 2009 N EUROPE. IEEE, Bremen, 2009; pp 1–7.

107. Langenkämper D, Zurowietz M, Schoening T, Nattkemper TW. BIIGLE 2.0 – Browsing and Annotating Large Marine Image Collections. Front Mar Sci 2017b; 4: 1–10.

108. Krizhevsky A, Sutskever I, Hinton GE. Imagenet classification with deep convolutional neural networks. In Krizhevsky A, Sutskever I, Hinton GE, editors. Advances in neural information processing systems. 2012; pp 1097–1105.

109. LeCun Y, Bengio Y, Hinton G. Deep learning. Nature. 2015; 521:436.

110. Woodall LC, Andradi-Brown DA, Brierley AS, Clark MR, Connelly D, Hall RA, Howell KL, Huvenne VAI, Linse K, Ross RE, Snelgrove P, Stefanoudis PV, Sutton TT, Taylor M, Thornton TF, Rogers AD. Multidisciplinary approach for generating globally consistent data on mesophotic, deep-pelagic, and bathyal biological communities. Oceanogr. 2018;31: 3.

111. Ebert DA, Stehmann MFW. Sharks, batoids, and chimaeras of the North Atlantic. FAO Species Catalogue for Fishery Purposes. No. 7. FAO, Rome; 2013.

112. Howell KL, Davies JS, van den Beld I. Deep-sea species image catalogue. University of Plymouth, Ifremer, NOAA. 2017; Available from: http://www.deepseacatalogue.fr/

113. Jones DOB, Gates AR, Curry RA, Thomson M, Pile A, Benfield M editors. SERPENT project. Media database archive. 2009; Available online: http://archive.serpentproject.com/

114. Rogacheva A, Gebruk A, Alt CH. Holothuroidea of the Charlie Gibbs Fracture Zone area, northern Mid-Atlantic Ridge. Mar Biol Res 2013;9: 587–623.

115. Oliveira F, Aguilar R, Monteiro P, Bentes L, Afonso CML, García S, Xavier JR, Ocana O, de Matos V, Tavares AM, Goncalves JMS. A photographic guide of the species of the Gorringe. Centro de Ciências do Mar/Oceana, Faro. 2017.

116. Kenchington E, Best M, Cogswell A, MacIsaac K, Murillo-Perez FJ, MacDonald B, Wareham V, Fuller SD, Jørgensbye HIØ, Sklya V, Thompson AB. Coral Identification Guide NAFO Area. NAFO Scientific Council Studies, Nova Scotia. 2009.

117. Best M, Kenchington E, MacIsaac K, Wareham VE, Fuller SD, Thompson AB Sponge Identification Guide NAFO Area. NAFO Scientific Council Studies, Nova Scotia. 2010; pp 43–50.

118. Kenchington E, Beazley L, Murillo FJ, Tompkins MacDonald G, Baker E. Coral, Sponge, and Other Vulnerable Marine Ecosystem Indicator Identification Guide, NAFO Area. NAFO Scientific Council Studies, Nova Scotia. 2015.

119. Packer D, Drohan A. Identification sheets for the common deep-sea corals off the Northeast and Mid-Atlantic US (v1.0). NOAA. 2013. Available from: https://www.nefsc.noaa.gov/fsb/training/NortheasternU.SDeepsea_Coral_Guide.pdf

120. NOAA Office of Ocean Exploration and Research Benthic Deepwater Animal Identification Guide. 2018. Available from: https://oceanexplorer.noaa.gov/okeanos/animal_guide/animal_guide.html

121. Serena F. Field identification guide to the sharks and rays of the Mediterranean and Black Sea. FAO Species Catalogue for Fishery Purposes. FAO, Rome; 2005.

122. Fourt M, Goujard A, Pérez T, Chevaldonné P. Guide de la faune profonde de la mer Méditerranée: Explorations des roches et canyons sous-marins des côtes françaises. Muséum national d’Histoire naturelle, Paris; 2017.

123. Xavier JR, Bo M. Deep-sea sponges of the Mediterranean Sea; 2017. Available from http://www.fao.org/3/a-i6945e.pdf

124. Bo M. Deep-sea corals of the Mediterranean Sea; 2017. Available from http://www.fao.org/3/a-i7256e.pdf

125. Alt CHS. On the benthic invertebrate megafauna at the Mid-Atlantic Ridge, in nity of the Charlie-Gibbs Fracture Zone. PhD Thesis, University of mpton. 2012.

